# Elephant Genomes Elucidate Disease Defenses and Other Traits

**DOI:** 10.1101/2020.05.29.124396

**Authors:** Marc Tollis, Elliott Ferris, Michael S. Campbell, Valerie K. Harris, Shawn M. Rupp, Tara M. Harrison, Wendy K. Kiso, Dennis L. Schmitt, Michael M. Garner, C. Athena Aktipis, Carlo C. Maley, Amy M. Boddy, Mark Yandell, Christopher Gregg, Joshua D. Schiffman, Lisa M. Abegglen

## Abstract

Disease susceptibility and resistance comprise important factors in conservation, particularly in elephants. To determine genetic mechanisms underlying disease resistance and other unique elephant traits, we estimated 862 and 1,017 potential regulatory elements in Asian and African elephants, respectively. These elements are significantly enriched in both species with differentially expressed genes involved in immunity pathways, including tumor-necrosis factor which plays a role in the response to elephant endotheliotropic herpesvirus (EEHV). Population genomics analyses indicate that amplified *TP53* retrogenes are maintained by purifying selection and may contribute to cancer resistance in elephants, including less malignancies in African vs. Asian elephants. Positive selection scans across elephant genomes revealed genes that may control iconic elephant traits such as tusk development, memory, and somatic maintenance. Our study supports the hypothesis that interspecies variation in gene regulation contributes to differential inflammatory responses leading to increased infectious disease and cancer susceptibility in Asian versus African elephants. Genomics can inform functional immunological studies which may improve both conservation for elephants and human therapies.

## Introduction

Elephants (family Elephantidae) first appeared on the planet ~5-10 million years ago (MYA) and three species roam today: the African bush elephant (*Loxodonta africana*), the African forest elephant. (*L. cyclotis*), and the Asian elephant (*Elephas maximus*). These species are the only surviving members of the once diverse proboscidean clade of afrotherian mammals^1^. Other elephantids such as straight-tusked elephants (genus *Paleoloxodon*) and mammoths (*Mammuthus*) went extinct around 34,000 and 4,300 years ago, respectively^2,3^. All extant elephant species are now threatened with extinction, largely due to poaching and habitat loss. Asian elephants are “endangered,” with only ~200 wild individuals in some countries^4^, and African elephants are “vulnerable” with only ~400,000 wild individuals after a decrease of ~100,000 individuals between 2007 and 2016^5,6^. It is imperative that we study the genetics of these amazing creatures and how this knowledge of their evolutionary history can contribute to their continued conservation.

Elephants share many charismatic traits such as prehensile trunks, ivory tusks, intelligence with long-term memory, and large body sizes^1^. Given their long lifespans of nearly 80 years for Asian^7^ and approximately 65 years for African elephants^8^, coupled with lengthy gestation periods of 22 months, disease defense has evolved as an important trait for elephants. Differences in disease susceptibility between species have urgent ramifications for elephant conservation^9^. In addition to poaching, Asian elephants are threatened by an acute hemorrhagic disease resulting from infection with elephant endotheliotropic herpesvirus (EEHV)^10,11^. While fatalities in African elephant calves from EEHV also have been reported, mortality rates are higher for Asian elephants suggesting a genetic component for increased EEHV lethality. Asian elephants also are more susceptible to tuberculosis (TB) infection (*Mycobacterium tuberculosis* and *M. bovis*)^9^ (Fisher’s Exact Test, P=2.52e-04; Chi-squared test, P=6.84e-04; Supplementary Fig. 1). Understanding the functional immunological and molecular basis of disease response in elephants may improve their conservation and medical care.

Another important disease for elephants is cancer, albeit for different reasons. Elephant cancer mortality rates are low compared to humans, despite the fact that cancer is a body size- and age-related disease^12^. A potential mechanism of cancer resistance for elephants is an enhanced apoptotic response of elephant cells to DNA damage associated with extensive amplification of retrogene copies of the tumor suppressor gene *TP53*^12–14^, yet there are still unknowns related to cancer in elephants. For instance, it is unclear if variation in *TP53* copy number contributes to differences in cancer defense between elephant species. Also, it is not known whether the observed differential responses to EEHV and TB between Asian and African elephants relate to cancer susceptibility. Detailed analyses of cancer prevalence and mortality in elephants may provide insight into how elephants evolved to handle disease. Here, we add to the knowledge of elephant cancer prevalence with data from zoos accredited by the Association of Zoos and Aquariums (AZA). AZA sets the standards for animal care and welfare in the United States (https://www.aza.org/about-us, last accessed September 2020). Every elephant in an AZA facility undergoes routine blood screens, full body examinations, and thorough necropsy upon death, making the likelihood of documenting elephant cancer high.

In addition to maintaining the health of elephants under human care to improve breeding and species survival plans, conservation efforts can benefit from genomic studies that identify genetic variants associated with traits such as disease defense^15^. However, the few elephant functional genomic studies currently available are limited to a small number of individuals and species^16–19^ and the genetic etiologies of most elephant traits are unknown. In our study, we analyze data from three living and two extinct species in comparative and population genomic frameworks in order to understand the genomic basis of elephant traits, including what drives different disease outcomes between species.

## Results

### Asian elephants suffer from higher rates of malignant cancers than African elephants

To estimate rates of neoplasia and malignancy in elephants, we collected and analyzed pathology data from 26 AZA-accredited zoos in the USA, which included diagnoses from 76 different elephants (n=35 African and n=41 Asian). We found that 5.71% of the African elephants and 41.46% of the Asian elephants were diagnosed with neoplasia (which included benign and malignant tumors) (Fisher’s Exact Test, P=3.78e-04; Chi-square test, P=8.95e-04) (Table 1, Supplementary Table 1). Sixty-nine percent of elephant tumors were benign, and 14.63% of Asian elephants were diagnosed with malignant tumors compared to zero in African elephants (Fisher’s Exact Test, P=0.028; Chi-squared test, P=0.053). In contrast, the lifetime risk of developing malignant cancer is 39.5% for humans^20^ and the lifetime risk of developing benign tumors is even higher, with 70%-80% of women developing uterine fibroids (leiomyomas) alone^21^. Asian elephants are also reported to have a high prevalence of uterine leiomyomas^22^, including seven in our dataset. Our results confirm that (1) malignant cancer rates in elephants are lower than in humans and (2) Asian elephants are diagnosed with both neoplasia and malignancies more often than African elephants in zoos.

**Table 1:** Cancer diagnoses and prevalence in African and Asian elephants.

| Neoplasia and Malignant Prevalence |  |  |  |  |  |
| --- | --- | --- | --- | --- | --- |
| Species | Total Individuals | Neoplasia | Neoplasia | Malignant | Malignant |
|  |  | Cases | % | Cases | % |
| Asian elephant | 41 | 17 | 41.46 | 6 | 14.63 |
| African elephant | 35 | 2 | 5.71 | 0 | 0 |
| Total elephants | 76 | 19 | 25 | 6 | 7.89 |
| Neoplasia Diagnoses in African/Asian Elephants |  |  |  |  |  |
| Species | Sex | Age | Lesion Type | Lesion Site | Malignant |
| African elephant | Female | 28 | Polyp | Vagina | No |
| African elephant | NA | NA | Adenoma | Parathyroid | No |
| Asian elephant | Female | 45 | Polyp | Vulva | No |
| Asian elephant | Female | 50;50 | Polyp;<br>Leiomyoma | Uterus;Uterus | No;No |
| Asian elephant | Female | 30;40 | Polyp; | Vagina; | No |
|  |  |  | Spindle Cell<br>Tumor | Leg |  |
| Asian elephant | Female | 39 | Leiomyoma | Uterus | No |
| Asian elephant | Female | 39 | Mast Cell Tumor | Abdomen | No |
| Asian elephant | Male | 35 | Papilloma | Trunk | No |
| Asian elephant | Female | 50 | Papilloma | Skin | No |
| Asian elephant | Female | 36 | Papilloma | Skin | No |
| Asian elephant | Female | 50 | Adenocarcinoma | Breast | Yes |
| Asian elephant | Female | 59;59 | Adenocarcinoma;<br>Leiomyoma | Uterus; Uterus | Yes; No |
| Asian elephant<br>(7) | NA | NA | Adenocarcinoma<br>(2); | Uterus (2); | Yes |
|  |  |  | Undifferentiated<br>Malignant<br>Neoplasm (1); | Uterus (1); |  |
|  |  |  | Leiomyosarcoma<br>(1); | Lung (1); |  |
|  |  |  | Sarcoma (1) | Liver (1) |  |
|  |  |  | Leiomyoma (4); | Uterus (4); | No |
|  |  |  | Leiomyoma (1); | Stomach (1); |  |
|  |  |  | Microadenoma (1) | Pituitary<br>Gland (1) |  |

### Elephant-specific accelerated genomic regions are enriched for immune pathways and correlate with species-specific gene expression patterns

To explore the genomic mechanisms governing disease response and other traits across elephant species, we sequenced and assembled the genome of an Asian elephant (“Icky”) born in Myanmar and currently under human care at 94.4X coverage with a final scaffold N50 of 2.77 Mb (GCA_014332765.1) (Supplementary Information, Supplementary Table 2, Supplementary Table 3, Supplementary Table 4). We also improved the African bush elephant genome assembly with Hi-C libraries (Supplementary Information, Supplementary Fig. 2, Supplementary Table 5). These genome assemblies were used to generate a whole genome alignment (WGA) with 10 other mammals, which we used to define accelerated genomic regions (ARs) unique to Asian and African elephants. We first defined 676,509 60 bp regions that were present in Asian and African elephants and conserved in the 10 background species with phastCons^23,24^ (conserved regions or CRs, Fig. 1a).

**Figure 1.**
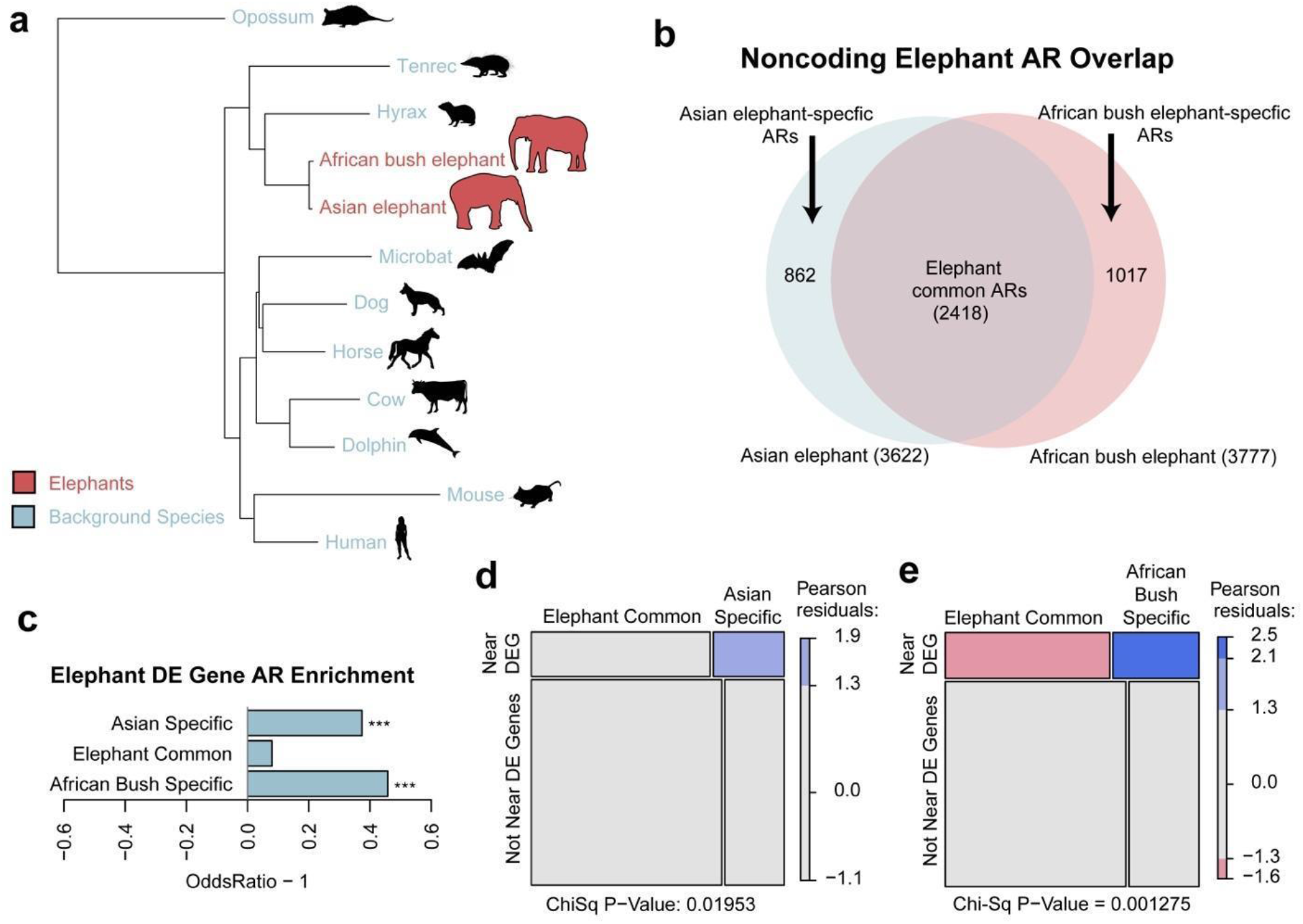
Elephant accelerated regions. Using a whole genome alignment of 12 mammals (a), we defined genomic regions that were accelerated (ARs) in two elephant species (red), yet conserved in the set of background species (blue). Branch lengths are given in terms of mean substitutions per site at fourfold degenerate sites (neutral model). Among the ARs detected in elephants, we found ARs common to both elephant species as well as ARs specific to either Asian or African bush elephants (b). Differentially expressed (DE) genes were much more likely to be found in Asian elephant-specific (Fisher’s Exact Test, P=2.05e-4) and African elephant-specific (P=8.30e-7) ARs than in common ARs (c). Species-specific ARs disproportionately overlap DE gene regulatory regions relative to the common ARs (Chi-squared test, p=0.019 and p=0.001 respectively) (d, e).

Asian and African elephants likely diverged ~5 MYA^25^, and since differences between closely related mammals are primarily due to changes in non-coding regulatory genomic regions^24,26,27^, we focused on the 376,899 CRs detected in non-coding regions. We tested these for accelerated substitution rates in elephants with phyloP^24,26^ and found 3,622 regions with significantly increased nucleotide substitution rates in the Asian elephant while 3,777 regions were accelerated in the African bush elephant (q-value < 0.10). We found 2,418 ARs shared between both species, with 862 Asian elephant-specific and 1,017 African bush elephant-specific ARs (Fig. 1b).

ARs common to Asian and African bush elephants were likely driven by changes pre-dating the evolutionary divergence of the two elephants, while Asian elephant- and African bush elephant-specific ARs may point to enhancers driving gene expression level changes that impact phenotypes distinguishing the two species. Using available African bush elephant and Asian elephant peripheral blood mononuclear cell (PBMC) RNA-Seq data^28,19^, we defined 5,034 differentially expressed (DE) elephant genes (false discovery rate or FDR < 0.01). Both Asian elephant- and African bush elephant-specific ARs were significantly enriched near DE genes relative to CRs (Fisher’s Exact Test, P=2.05e-4, P=8.30e-7, respectively). Meanwhile, the 2,418 ARs common to both elephants were not significantly enriched near DE genes (Fig. 1c). This pattern remained robust with subsets of increasingly significantly DE genes based on adjusted p-values (Supplementary Fig. 3). Asian elephant- and African bush elephant-specific Ars disproportionately overlapped DE gene regulatory regions relative to the common ARs (Chi-squared test, P=0.019 and P=0.001 respectively; Fig. 1d, Fig. 1e), suggesting that some ARs reflect changes in regulatory regions that alter gene expression patterns in elephant PBMCs.

We functionally annotated AR contributions to African and Asian elephant species differences by testing the elephant species-specific and common ARs for Biological Process (BP) gene ontology (GO) term enrichments (Supplementary Data 1). Based on a likelihood ratio test that compared general linearized models (see Materials and Methods), 605 out of 607 (99.6%) of GO terms were uniquely enriched in the elephant ARs in contrast to ARs found in other mammalian lineages^19^. This suggests that the enrichment of GO terms in the elephant ARs are significant in elephants in contrast to other mammals. Of 18,056 BP GO terms, 252 were significantly enriched in Asian elephant specific ARs and 275 were enriched in African elephant specific ARs (q-value < 0.05). Many of the GO terms related to the immune system in both elephant species (Fig. 2; Supplementary Fig. 4; Supplementary Fig. 5; Supplementary Data). The broad term ‘immune system process’ was 4.5 and 2.8 fold enriched with Asian and African elephant-specific ARs (q-value = 4.87e-12 and 7.84e-06, respectively), but not significantly enriched with elephant common ARs. Our results suggest (1) many of the species-specific ARs alter gene expression patterns and transcription factor binding networks that eventuate differences in immune function, and (2) Asian-elephant ARs are more enriched in immune pathways than African elephant-specific ARs in terms of both fold-enrichment and statistical significance.

**Figure 2.**
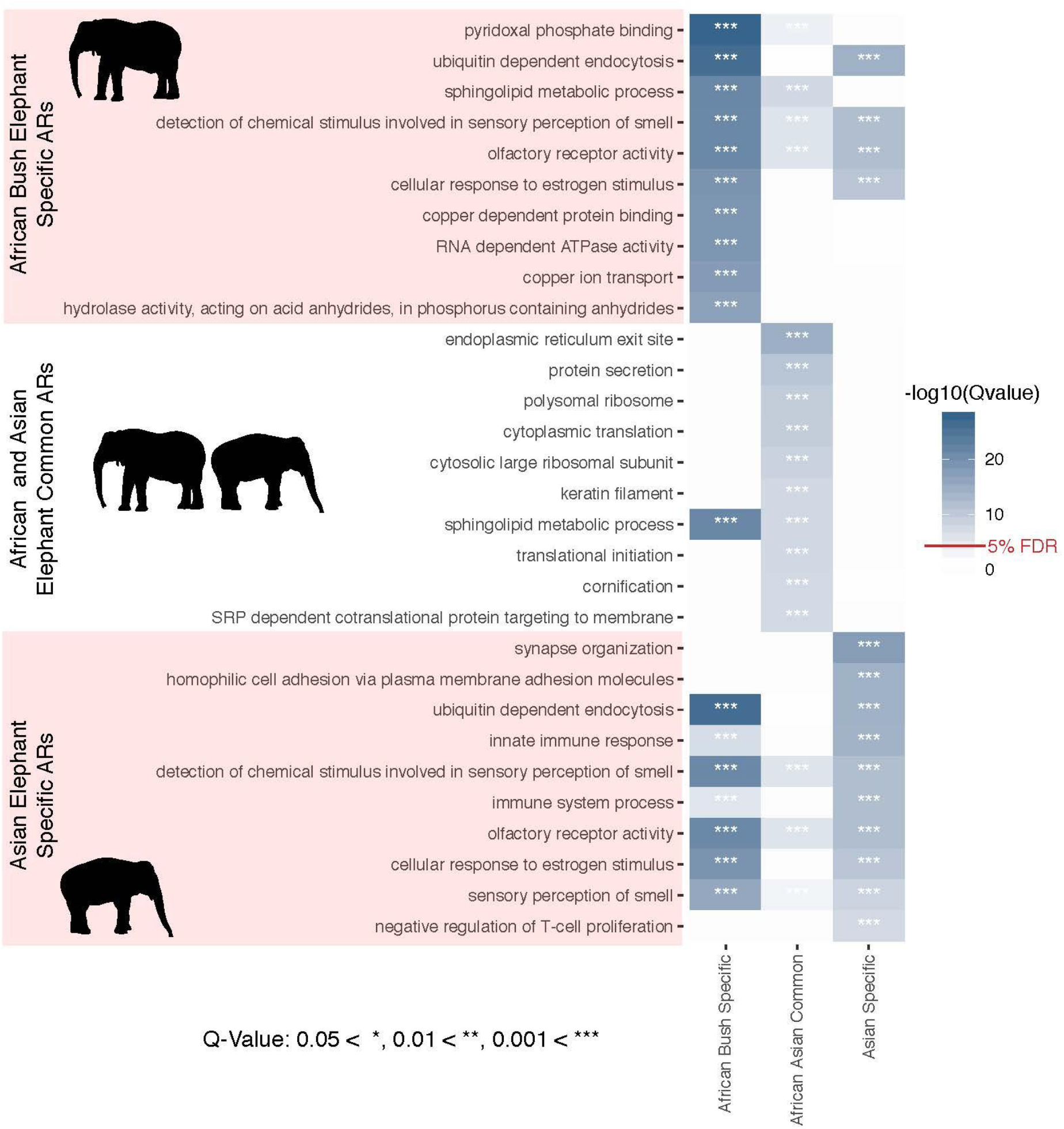
Top 10 Biological Process gene ontology (GO) terms most significantly enriched with elephant accelerated genomic regions (ARs). The 10 most significantly enriched GO terms in terms of −log10(q-value) for each set of ARs (African elephant-specific, African and Asian elephant common, Asian elephant-specific), and their overlap. “Innate immune response” and “immune system process” are in the top 10 most significantly enriched GO terms for Asian elephant-specific ARs, are significantly enriched in African elephant specific ARs but not in the top 10, and are not significantly enriched in the common ARs. “Negative regulation of T-cell proliferation” was only in top 10 significantly enriched GO terms for the Asian elephant-specific ARs.

We found 109 GO terms significantly enriched with elephant common ARs (q-value < 0.05, Supplementary Fig. 6, Supplementary Data), many of which were related to cancer, including “sphingolipid metabolic process” which was in the top 10 most significantly enriched GO terms for both elephant common (5.7 fold enrichment, q-value = 4.69e-08) and African elephant-specific (17.3 fold enrichment, q-value = 4.18e-22) ARs (Fig. 2). Sphingolipid metabolites mediate the signalling cascades involved in apoptosis^29^, necrosis^29^, senescence^30^, and inflammation^31^. We found 2.9 and 3.6 fold enrichments for ‘tumor necrosis factor (TNF)-mediated signaling pathway’ (q-value=4.75e-04) and ‘positive regulation of TNF production’ (q-value = 1.75e-03) in the common ARs, and a 21.5 fold enrichment of “negative regulation of TNF secretion” in African elephant-specific ARs (q-value = 5.01e-04). TNF is a cytokine involved in cell differentiation and death that can induce the necrosis of cancer cells^34^. The upregulation of TNF-alpha has been associated with increased apoptosis in EEHV-infected Asian elephant PBMCs as well^11^.

In a check for reproducibility, we found that the number of African elephant-specific ARs assigned to each gene was correlated with previous studies^19^ (R = 0.82). The gene most enriched with previously defined non-coding African elephant ARs was the DNA repair gene *FANCL* (7.3 fold enrichment; q-value = 2.16e-56), which mediates the E3 ligase activity of the Fanconi anemia core complex, a master regulator of DNA repair ^32^. We found that *FANCL* was the third most significantly enriched gene in both African and Asian elephant ARs relative to CRs with 4.6 and 4.9 fold enrichments (q-value = 1.27e-14 and 4.46e-16). Of 50 African elephant ARs and 51 Asian elephant ARs assigned to *FANCL*, 43 are common to both elephant species suggesting their acceleration predates African-Asian elephant divergence. These results suggest non-coding cis-regulatory elements have regulated cancer resistance adaptations throughout elephant evolution, particularly in the ancestor of modern elephants and the lineage leading to the African bush elephant.

### Evolution of *TP53* and its retrogenes in elephant genomes

The enhanced apoptotic response to DNA damage in elephant cells correlates with the expansion of ~20 copies of the tumor suppressor gene *TP53* in elephant genomes^12–14^, and we wanted to (1) understand the evolutionary origins of *TP53* gene duplications in Asian and African bush elephants and (2) determine if *TP53* copy number is related to body size evolution in elephants. We annotated *TP53* homologs in 44 mammalian genomes including Icky the Asian elephant, an additional genome assembly of an Asian elephant^33–35^ (“Methai”, born in Thailand and living at the Houston Zoo, assembly available at www.dnazoo.org, last accessed September 2020), and the African bush elephant assembly presented in this study (Supplementary Table 6), and incorporated them in a molecular clock analysis. We estimated that *TP53* retrogene copies originated in the paneungulate ancestor of manatees and elephants ~55-60 million years ago (MYA) (41.3–75.2 95% highest posterior density or HPD) (Fig. 3a, Supplementary Fig. 7). A subsequent TP53 expansion began ~45 MYA (30.7–60.1 95% HPD) in a common ancestor of Asian and African elephants, and continued throughout elephantid evolution. We estimated 19 copies of TP53 in the African bush elephant genome assembly, and 9-11 TP53 copies in the Asian elephant genome based on the two assemblies for the species.

**Figure 3.**
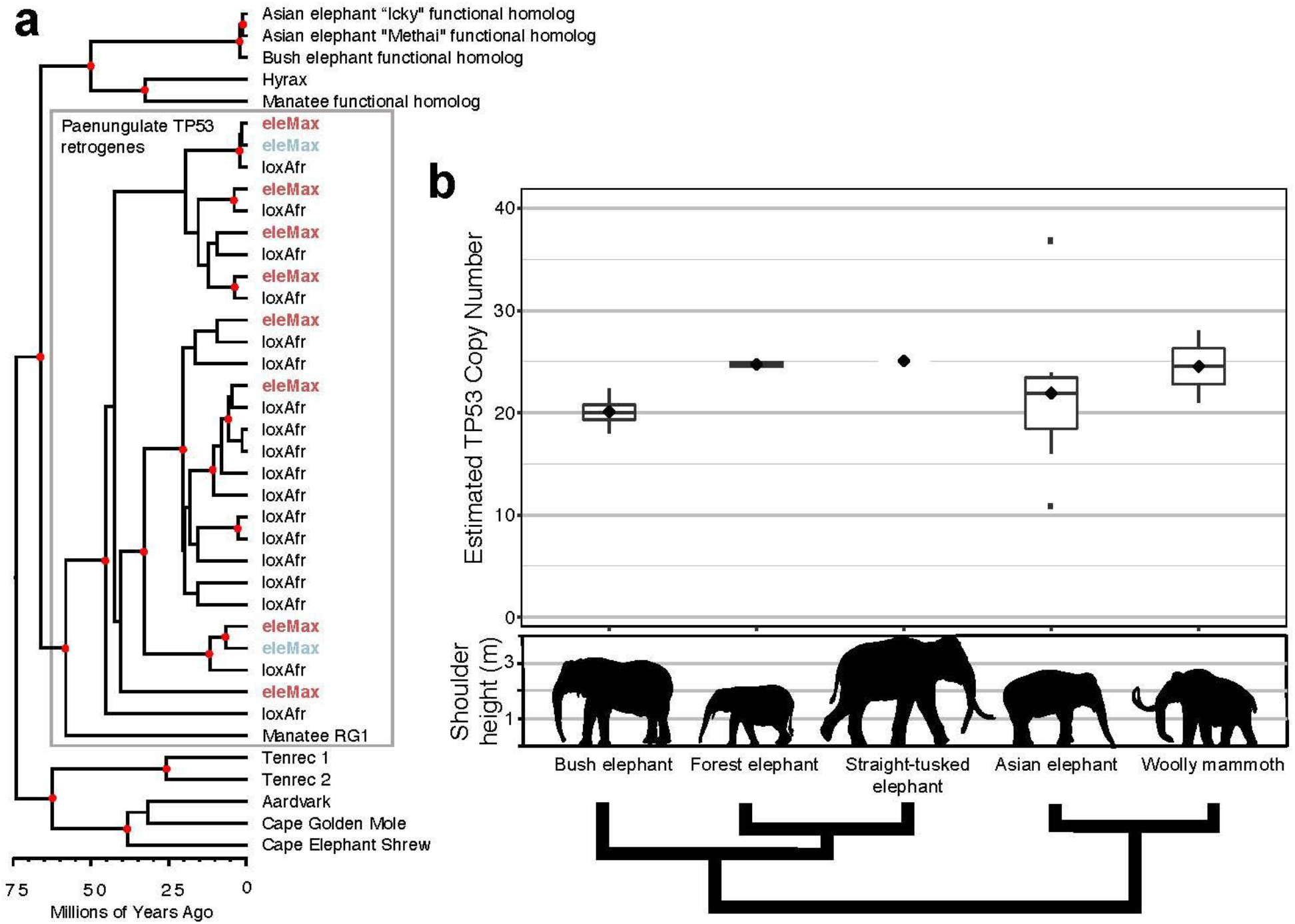
Evolution of *TP53* in elephants and other afrotherians. (a) Phylogeny of *TP53* sequences extracted from afrotherian genomes. *TP53* retrogenes (Manatee RG1, eleMax, and loxAfr) appeared early in the evolution of paenungulates ~55–60 million years ago (MYA), followed by subsequent amplification in the elephant lineage ~45 MYA. Red dots indicate estimated nodes with posterior probability ≥90%. Red eleMax indicates *TP53* retrogene sequences extracted from the “Methai” Asian elephant assembly, and blue eleMax indicates *TP53* retrogene sequences extracted from the Asian elephant assembly “Icky” presented in this study. (b) *TP53* copy number estimates based on read counts from three living elephants (African bush elephant (n=4, minimum 18, maximum 22.4, median 20.1, 25th percentile 19.3, 75th percentile 20.8), forest elephant n=2, minimum 24.3, maximum 25.2, median 24.7, 25^th^ percentile 24.5, 75th percentile 25) and Asian elephant (n=7, minimum 10.9, maximum 36.8, median 21.9, 25th percentile 18.5, 75th percentile 23.4)) and two extinct (straight-tusked (n=1, 25.1) and woolly mammoth (n=2, minimum 21, maximum 28, median 24.51, 25th percentile 22.8, 75th percentile 26.3)) elephant species. Shoulder height estimates from Larramendi et al. (2015). Phylogeny is schematic only and represents relationships from Palkopuolou et al. (2018).

We mapped whole genome shotgun data from multiple individuals belonging to three living and two extinct elephant species (Supplementary Table 7) to the bush elephant genome annotation (loxAfr3) and used normalized read counts to estimate *TP53* copy numbers in elephant genomes (Figure 3b; Supplementary Table 8). Based on read depth, African bush elephants (n=4) have on average ~19–23 *TP53* copies in their genomes, and Asian elephant genomes (n=7) contain as few as 10 *TP53* copies, or as many as 37, but without these outliers average ~19–22 *TP53* copies in their genomes. These estimates are similar to previous estimates of *TP53* copy numbers for bush and Asian elephants based on smaller numbers of individuals^12–14^. We estimated ~21–24 *TP53* copy number variants in forest elephant genomes (n=2). The woolly mammoths (n=2) were estimated to have between 19 and 28 *TP53* copies in their genomes, which was slightly higher than previous estimates^14^. Meanwhile, the straight-tusked elephant genome contained ~22-25 *TP53* copies.

The number of *TP53* copies estimated in the genomes of Asian elephants differed based on the method used. For instance, we validated two *TP53* retrogenes in Icky’s genome assembly, which phylogenetically clustered closely with two of the nine retrogenes validated in Methai’s genome assembly (Fig. 3a). Taken together, based on the reciprocal BLAT searches of both assemblies we estimated 9-11 *TP53* copies in the Asian elephant genome. However, based on normalized read counts, we estimated 10-37 TP53 copies (Fig. 3b), which is more consistent with previous studies^12,14^. The lower estimates we obtained from the Asian elephant genome assemblies may be due to poorly resolved repetitive regions which hamper graph-based *de novo* genome assemblers^33^. Subsequent refinement of Asian elephant genome assemblies using long read sequencing may better resolve these regions. In the meantime, our results suggest that copy number estimates based on read depth are useful approximations for approaches validated from genomic DNA.

### A bottleneck for bush elephants and a deep divergence for forest elephants

Our next goals were to (1) determine if *TP53* paralogs segregate as functional alleles in elephant populations and (2) estimate regions of adaptive evolution (positive selection) that may control phenotypes in modern elephants. To do this, we utilized the aligned genomic sequences from the living elephant species to call variants with freebayes v1.3.1-12^36^, genotyping 41,296,555 biallelic single nucleotide polymorphisms (SNPs), averaging one SNP every 77 bases and with a genome-wide transition-transversion ratio of 2.46. Altogether, we annotated 290,965 exonic, 11,245,343 intronic, and 32,512,650 intergenic SNPs across the 13 elephant genomes.

To establish the neutral background against which to compare putative adaptively evolving regions of the genome, we assessed the demographic history of each elephant species with summary population genomics statistics. Among the three living species of elephants, bush elephants averaged the lowest nucleotide diversity (0.0008, s.d. 0.001), followed by Asian elephants (0.0011, s.d. 0.001) and forest elephants (0.002, s.d. 0.002) (Fig. 4a). The distribution of Tajima’s D in 10 Kb genomic bins calculated for bush elephant revealed an excess of negative values relative to other elephant species (Fig. 4b). We also found a larger proportion of heterozygous sites (0.18 and 0.21) in forest elephant genomes compared to all other elephants (Fig. 4c), consistent with the deep genomic divergence reported in this species^25,37^.

**Figure 4.**
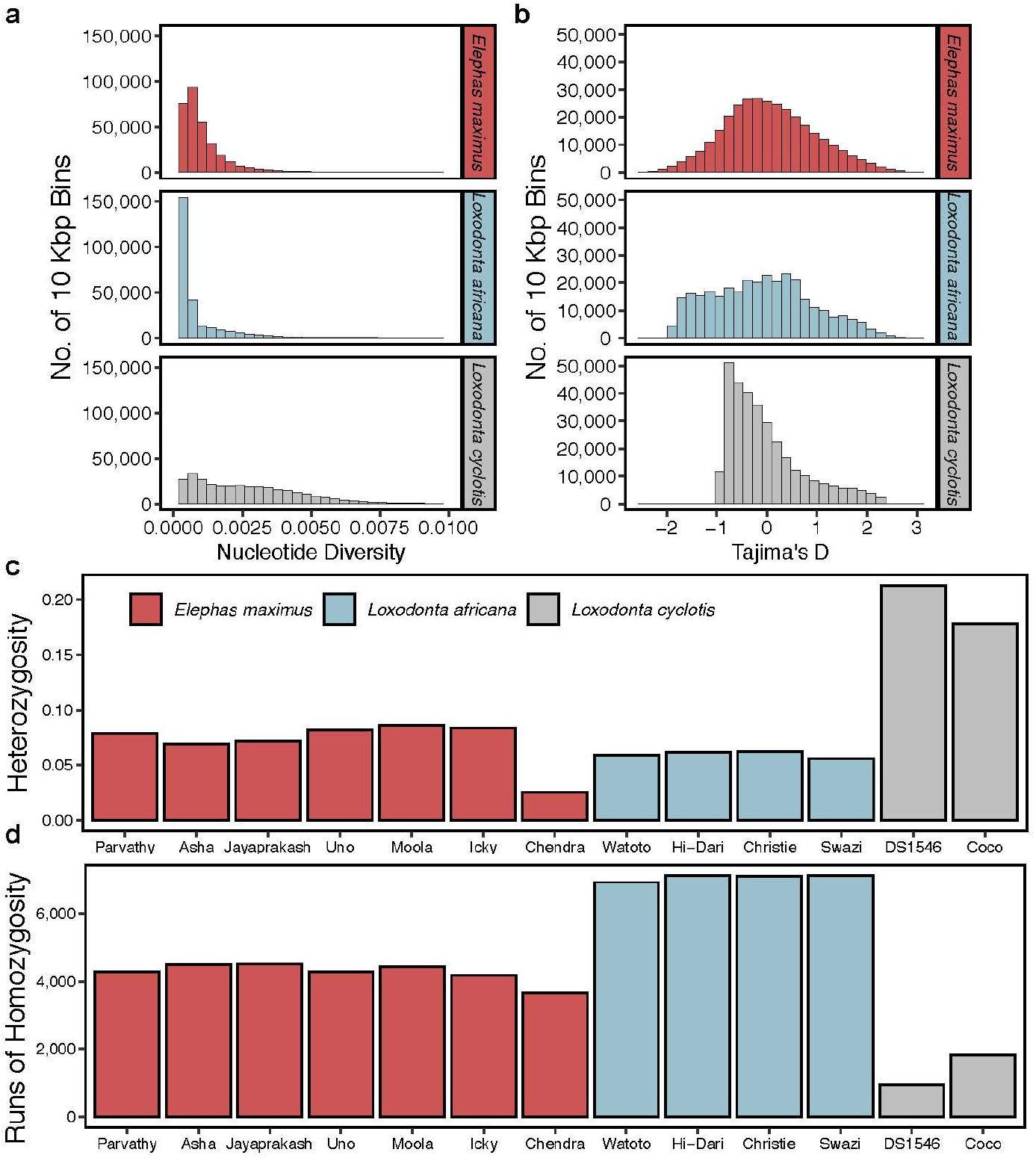
Summary population genomics statistics for three living elephant species. We analyzed 10 Kbp bins for (a) nucleotide diversity and (b) Tajima’s D for Asian (*Elephas maximus*), bush (*Loxodonta africana*), and forest (*L. cyclotis*) elephants. (c) Per-individual heterozygosity for Asian, bush, and forest elephants. (d) Number of detected runs of homozygosity for each individual elephant.

After identifying runs of homozygosity (RoH) in each elephant (Fig. 4d, Supplementary Fig. 8a), we estimated the average inbreeding coefficient (*F_RoH_*) to be higher in bush elephants (2.04%) than Asian (1.64%) and forest (0.61%) elephants (Supplementary Fig. 8b). Taken together, the excess of low frequency polymorphisms, RoH, and *F_RoH_* suggest a strong population bottleneck in bush elephants.. The Asian elephant from Borneo contained the smallest number of heterozygous genotyped sites among all elephants (0.03), and the highest *F_RoH_* (5.39%), consistent with recent analyses showing that the Borneo subpopulation of *E. maximus* is genetically isolated^25^.

### Genetic variation in *TP53* copy number variants suggests maintenance of some by purifying selection

We found a high degree of sequence conservation in the *TP53* paralogs both within and between the three living elephants (Supplementary Table 9). For instance, the proportion of polymorphic sites in putatively neutrally evolving ancestral repeats was 0.013, but across 12 annotated *TP53* paralogs was 0.004. Despite the deep genomic divergence of forest elephants, we found very little genetic variation in *TP53* paralogs for the species, with just a single segregating site in three of the retrogenes. Across all species, we collected zero nonsynonymous SNPs for the functional homolog (ENSLAFG00000007483), consistent with strong purifying selection on this gene and ENSLAFG00000028299, or “retrogene 9”, whose protein expression is stabilized by DNA damage in human cells^12^.

We annotated variants in 12 *TP53* paralogs based on the bush elephant genome annotation and found few variants affecting gene function (Supplementary Table 10), consisting of mostly missense mutations. There were no variants of high or moderate impact on gene function annotated in the functional homolog (ENSLAFG00000007483), with the majority of variants occurring downstream, in introns, or upstream of the gene. The high degree of sequence conservation across three species of elephant, and in particular the lack of variants with functional effect, especially in “retrogene 9,” suggests that at least some *TP53* retrogenes are being maintained by purifying selection.

### An elephant never forgets: positive selection in living elephants

To assess the impact of natural selection across elephant genomes and its impact on phenotypes, we scanned the genomes of the three extant species for positive selection using SweeD v3.3.1^38,39^, hypothesizing that genetic pathways controlling elephant traits would be subjected to selective sweeps. This yielded 24,394 selectively swept outlier regions meeting our statistical thresholds based on neutral expectations (see Materials and Methods) in Asian elephants, which comprise ~0.07% of the genome and overlapped with 1,611 gene annotations. Out of the 41,204 regions experiencing putative selective sweeps in bush elephants (~1.3% of the genome), we detected 2,882 protein-coding genes. We estimated 4,099 protein-coding genes involved in the 51,249 regions involved in putative selective sweeps in forest elephants (~1.6% of the genome).

We found 242 protein-coding genes that overlapped regions with evidence of positive selection and were shared in all three of the living elephant species, which are enriched in BP GO terms that shed light on the genomic mechanisms controlling many iconic elephant traits. For instance, the most significantly enriched BPs (in terms of fold enrichment) were “dendrite self-avoidance” (27-fold enrichment, FDR=0.02), “ionotropic glutamate receptor signaling pathway” (17-fold enrichment, FDR=0.02), and “regulation of NMDA receptor activity” (14-fold enrichment, FDR=0.03). Many significantly enriched GO terms clustered semantically with “trans-synaptic signaling” (Fig. 5a). We also found significant protein interactions among outlier genes (75 observed edges versus 45 expected; enrichment P=3.68E-5), including an enrichment of genes associated with the glutamatergic synapse pathway (7 of 98 genes, FDR=0.015). Glutamate is the major excitatory neurotransmitter for mammalian nerve cells, mediating excitatory synaptic transmission^40^. These results suggest strong selection in elephants on pathways involved in memory, learning, and the formation of neural networks. The recognition, storing and retrieving of information in the human brain occurs in the temporal lobe, and the temporal lobes of elephants’ brains are relatively larger than those of humans, as well as denser and more convoluted^41^. Retrieving information is likely crucial for elephants to find resources across vast and complicated landscapes^1^.

**Figure 5.**
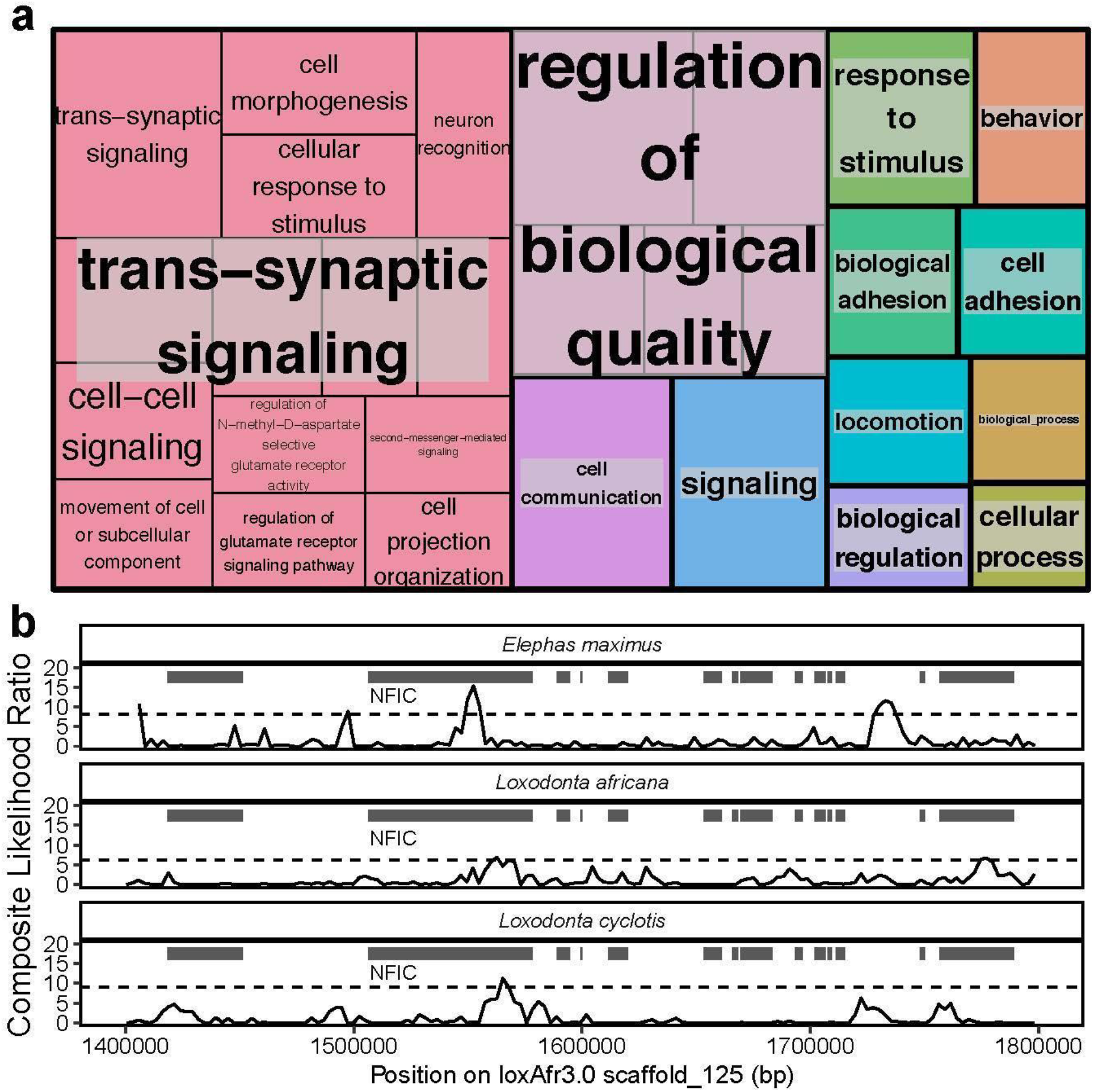
Selective sweeps in three living elephant species. (a) TreeMap from REVIGO representing semantic clustering of gene ontology biological process terms with a Benjamini-Hochberg false discovery rate of 5% that are associated with genes overlapping selective sweeps common to *Elephas maximus*, *Loxodonta africana*, and *L. cyclotis.* Rectangles represent clusters, and larger rectangles indicate semantically related clusters. Larger rectangle sizes reflect smaller corrected p-values from the GO term enrichment. (b) Composite likelihood ratio values in the *NFIC* region of a genomic scaffold (loxAfr3.0) calculated with SweeD in three elephant species. Gene annotations are represented by dark rectangles; the NFIC gene is indicated. Dashed lines represent p-value threshold of 0.0001.

Our genomic results suggest that positive selection has acted on genes involved in elephant tusk formation. Unlike most tusked mammals which feature elongated canines, the elephant tusk is a highly modified upper incisor^1^. We found that the most significantly enriched mouse phenotypes among genes overlapping selective sweeps were “abnormal upper incisor morphology” (FDR=0.001) and “long upper incisors” (FDR=0.003). These included two genes: *ANTXR1* and *NFIC*, (Figure 5b) which are involved in tooth development in humans^42–44^ and mice^45^, respectively.

Other significant gene ontology terms from outlier regions in our selective sweep analysis were related to cancer, including cell adhesion (9-fold enrichment, FDR=0.007), cell-cell signaling (3-fold enrichment, FDR=0.01), and cell communication (2-fold enrichment, FDR=0.0001). Significant protein-protein interactions were found associated with EGF-like domain (UniProt keyword enrichment, 13 out of 209 genes, FDR=4.2e-04; and INTERPRO protein domain enrichment, 13 out of 191 genes, FDR=2.6e-04). The EGF-like domain contains repeats which bind to apoptotic cells and play a key role in their clearance^46^. Our selective sweep results are consistent with those from the AR analysis and suggest ongoing selection in pathways involved with somatic maintenance and in particular apoptosis, a possible key mechanism for cancer suppression in elephants.

## Discussion

Our study of elephant genomes expands the knowledge of elephant evolution, highlighting differences and similarities between species. Elephant tumors tend to be benign with strong genetic defenses to prevent malignant transformation. Asian elephants reported in our study develop benign tumors and malignant cancer at higher rates than African elephants, and therefore may benefit from increased monitoring for tumors. Even though our data originates from captive elephants, these differences most likely reflect true genetic differences as the AZA has a Species Survival Plan (SSP) (https://www.aza.org/species-survival-plan-programs) for elephants that maximizes genetic diversity via the careful selection of mate pairs and studbook documentation^47,48^. Together with the fact that many elephants in zoos are wild born, it is likely that wild Asian elephants share increased susceptibility to neoplasia with the same observed genetic variation we report in our study.

While collecting cancer prevalence data in wild elephants is challenging due to decomposition and predator consumption, future data from wild elephants and genomic analysis of benign vs. malignant tumors will be crucial to further understand the evolutionary basis of differences in cancer risk between elephant species. This information could benefit the survival of individual elephants and assist with selecting the best treatment interventions when the rare elephant tumor is diagnosed in captivity or in the wild. More than half of the elephant tumors reported here were found in reproductive organs (Table 1). Even benign reproductive tumors can affect reproduction and conservation, therefore future studies to understand their impact and to develop preventative and treatment measures are needed.

While previous studies suggest that *TP53* copy number increased with body mass during proboscidean evolution as a response to increased cancer risk^14^, we estimated some of the highest *TP53* copy numbers in the smallest elephants. Based on available sequence data, we estimated ~19–21 *TP53* copy number variants in the ~44,800 year old woolly mammoth genome from Oimyakon, Russia, but found that the much more recent ~4,300-year-old Wrangel Island mammoth had ~1.3X this number of *TP53* copies in its genome. These findings are consistent with the demographic decline of the last woolly mammoths on Wrangel Island^18^, which by 12,000 years before present shrunk in body size by ~30% relative to more ancient mammoths elsewhere^4^. The estimated *TP53* copy number increase in the Wrangel Island mammoth may be related to the random fixation of retrogenes in the population rather than selection acting on body size.

We estimated ~21–24 *TP53* copy number variants in forest elephant genomes, greater than our estimates for bush elephants despite the smaller body size of forest elephants. Meanwhile, we estimated ~23–25 *TP53* copy number variants in the genome of the straight-tusked elephant, which at ~13,000kg may have been the largest land mammal to have ever lived (Fig. 3b)^49^. Recent genomic evidence suggests that forest elephants are more closely related to straight-tusked elephants than to bush elephants (Fig. 3b)^25,50^, with extensive gene flow occurring between forest and straight-tusked elephants, as well as between straight-tusked elephants and mammoths^25^. Thus, the possibility exists that, as in the Wrangel Island mammoth, the higher estimated *TP53* copy number in forest elephants relative to bush elephants may not be related to modern differences in body mass between species (and possible protection from increased cancer risk), but instead may be due to complicated evolutionary and demographic histories which include migration that can dramatically affect the dynamics of repetitive genomic elements such as retrogenes^51^. Nevertheless, we still find that genetic variation at some *TP53* retrogenes is tightly conserved in populations of all living elephant species. This adds at least some evidence to the functionality of *TP53* retrogenes. We suggest that there may be a core set of *TP53* retrogenes that confer the bulk of cancer suppression in elephants.

Our results support the idea that regulatory elements play a role in the increased infectious susceptibility with inflammatory response of Asian versus African elephants, particularly the mediation of the TNF cytokine. Asian elephant calves are much more susceptible than African elephant calves to cytokine storm triggered by EEHV infection^52^. Compared to African elephants, we found that Asian elephant ARs are enriched for BP GO terms “interleukin-1 beta (IL-1β) production” (q-value=0.036), “interleukin-18 (IL-18) production” (q-value=0.00073), and “neutrophil activation involved in immune response” (q-value=2.44e-05) (Supplementary Data). IL-1β, IL-18 and neutrophils, combined with TNF-alpha, are potent mediators of innate immunity. Uncontrolled activation of these factors leads to immune-induced disease pathogenesis through excessive inflammation. In humans and other mammals, neutrophil activation directly contributes to tissue damage through the release of neutrophil extracellular traps (NETs)^53,54^. Functional studies to compare cytokine secretion and NET release in response to infectious agents are ongoing and could confirm that genetic differences in innate immunity contribute to differences in disease susceptibility and outcomes between Asian and African elephants. Our study provides an example of how genomics can inform functional immunological and molecular studies, which may lead to improved conservation and medical care for elephants. This type of genetic information could provide important evolutionary insights to one day be translated into human patients with infection or cancer.

## Materials and Methods

### Cancer Data Collection

Pathology and necropsy records were collected with consent from 26 zoological institutions across the United States over the span of 26 years. All participating institutions were de-identified and anonymized. Data was collected with permissions from Buffalo Zoo, Dallas Zoo, El Paso Zoo, Fort Worth Zoo, Gladys Porter Zoo, Greenville Zoo, Jacksonville Zoo and Gardens, Louisville Zoological Garden, Oakland Zoo, Oklahoma City Zoo and Botanical Garden, Omaha’s Henry Doorly Zoo and Aquarium, The Phoenix Zoo, Point Defiance Zoo and Aquarium, San Antonio Zoological Society, Santa Barbara Zoological Gardens, Sedgwick County Zoo, Seneca Park Zoo, Toledo Zoological Gardens, Utah’s Hogle Zoo, Woodland Park Zoo, Zoo Atlanta, Zoo Miami and three other anonymous zoos. Neoplasia was diagnosed by board certified veterinary pathologists and cancers were identified from pathological services at Northwest ZooPath, Monroe, WA. Published pathology data from San Diego Zoo was also included^55^. Neoplasia prevalence was estimated by the number of elephants that were diagnosed with tumors (benign or malignant) in respect to all elephants documented within our database.

### De novo assembly and annotation of the Asian elephant genome

A whole blood sample was drawn in an EDTA tube from the Asian elephant (“Icky”, North American studbook number 199) from the Ringling Bros. Center for Elephant Conservation, and DNA libraries were constructed and sequenced at the University of Utah Genomics Core Facility. Paired-end DNA libraries were constructed with the TruSeq Library Prep kit for a target insert size of 200 bp, and mate-paired libraries were constructed for target sizes of 3 kb, 5 kb, 8 kb, and 10 kb using the Nextera Mate Pair Library kit. Genomic DNA was sequenced on an Illumina HiSeq2500. Raw reads were trimmed to remove nucleotide biases, adapters and a quality score cutoff of 30 with Trimmomatic v0.33^56^ and SeqClean^57^. Genome assembly was carried out using ALLPATHS-LG^58,59^. The expected gene content was assessed by searching for 4,104 mammalian single-copy orthologs (mammalia_odb9) using BUSCO v3.0.2^60^. We annotated and masked repeats in the resulting assembly using both the *de novo* method implemented in RepeatModeler v1.0.11^61^ and a database method using RepeatMasker v4.07^62^ with a library of known mammalian repeats from RepBase^63^. Modeled repeats were used in a BLAST search against Swiss-Prot^64^ to identify and remove false positives. We then generated gene models for the Asian elephant assembly using MAKER2^65^, which incorporated (1) homology to publicly available proteomes of cow, human, mouse, and all mammalian entries in UniProtKB/Swiss-Prot, and (2) ab initio gene predictions based on species-specific gene models in SNAP (11/29/2013 release)^66^, species-specific and human gene models in Augustus v3.0.2^67^, and EvidenceModeler^68^. Final gene calls were functionally annotated by using InterProScan^69^ to identify protein domains and a blastp search of all annotated proteins to UniProt proteins to assign putative orthologs with an e-value cutoff of 1e-6.

### Tissue collection, DNA extraction, and genome sequencing of African bush elephants

The African bush elephant assembly was improved with the addition of Hi-C sequencing libraries. First, a whole blood sample was drawn (in an EDTA tube) from a wild-born female named Swazi (animal ID: KB13542, North American studbook number 532) at the San Diego Zoo Safari Park in Escondido, CA. We selected this individual because her genome was originally sequenced by the Broad Institute^25^. Three Hi-C libraries were constructed and sequenced to ~38X genome coverage and used for scaffolding with HiRise^70^ at Dovetail Genomics in Santa Cruz, CA, with the most recent version of the African bush elephant assembly (loxAfr4.0) as an input. DNA was extracted from fresh frozen subcutaneous skin necropsy tissue samples from an African bush elephant named Hi-Dari (animal ID 00003, North American studbook number 33) at the Hogle Zoo in Salt Lake City, UT using a ThermoFisher PureLink Genomic Mini DNA Kit at the University of Utah. Two pieces of tissue were digested and extracted separately and pooled followed extraction. DNA concentration was measured by PicoGreen (8.66ng/ul) with a total volume of 200ul in 10mM pH8.0. DNA sequencing libraries were generated using the Illumina TruSeq Library Prep Kit for a 350 bp mean insert size, and sequenced on two lanes the Illumina HiSeq2500 platform at Huntsman Cancer Institute’s High-Throughput Genomics Core (Salt Lake City, UT).

### TP53 evolution in African and Asian elephants

To determine *TP53* copy numbers and evolutionary patterns across placental mammals, we exported the *TP53* human peptide from Ensembl (July 2019), and used it as a query in reciprocal BLAT searches^71^ of 44 mammalian genome assemblies (Supplementary Table 6), validated with a BLASTX query of the human peptide database on NCBI in order to ensure the top hit was human TP53 with ≥70% protein identity, following Tollis et al. (2020)^72^. Accepted nucleotide sequences were aligned with MAFFT^73^, and we weighted and filtered out unreliable columns in the alignment with GUIDANCE2^74^ using 100 bootstrap replicates. We reconstructed the phylogeny of all aligned mammalian TP53 homologs and estimated their divergence times in a Bayesian framework with BEAST 2.5^75^ using the HKY substitution model, a relaxed lognormal molecular clock model, and a Yule Model tree prior. We used a normal prior distribution for the age of Eutheria (offset to 105 million years or MYA with the 2.5% quantile at 101 MYA and the 97.5% quantile at 109 MYA) and lognormal prior distributions for the following node calibrations from the fossil record^61^: Boreoeutheria (offset the minimum age to 61.6 MYA –164 MYA and the 97.5% quantile to 170 MYA), Euarchontoglires (56 MYA – 164 MYA), Primates (56 MYA – 66 MYA), Laurasiatheria (61.6 MYA – 164 MYA) and Afrotheria (56 MYA – 164 MYA). We monitored proper MCMC mixing with Tracer v1.7.1^76^ to ensure an effective sampling size of at least 200 from the posterior distributions of each parameter and ran two independent chains. The final MCMC chain was run for 100,000,000 generations, and we logged parameter samples every 10,000 generations to collect a total of 10,000 samples from the posterior distribution. We then collected 10,000 of the resulting trees, ignored the first 10% as burn-in, and calculated the maximum clade credibility tree using TreeAnnotator.

### Detection of accelerated regions in African and Asian elephant genomes

We generated a multiple alignment (whole genome alignment or WGA) of 12 mammalian genome assemblies. First, we downloaded publicly available pairwise syntenic alignments of opossum (monDom5), mouse (mm10), dolphin (turTru1), cow (bosTau7), dog (canFam3), horse (equCab2), microbat (myoLuc1), tenrec (echTel2), and hyrax (proCap1) to the human reference (hg19) from the UCSC Genome Browser^77^. We also computed two additional *de novo* pairwise syntenic alignments with the human genome as a target and the two elephant genome assemblies reported here as queries using local alignments from LASTZ v1.02^78^ using the following options from the UCSC Genome Browser for mammalian genome alignments: -- hspthresh 2200 --inner 2000 --ydrop 3400 --gappedthresh 10000 --scores HOXD70, followed by chaining to form gapless blocks and netting to rank the highest scoring chains^79^. We then constructed a multiple sequence alignment with MULTIZ v11.2^80^ with human as the reference species.

To define elephant accelerated regions (ARs), we used functions from the R package rphast v1.6^24^. We used phyloFit with the substitution model ‘REV’ to estimate a neutral model based on autosomal fourfold degenerate sites from the WGA. We then used phastCons to define 60 bp autosomal regions conserved in the 10 non-elephant species in the WGA with the following options: expected.length = 45, target.coverage = 0.3, rho = 0.31. We further selected regions with aligned sequence for both African and Asian elephants that have aligned sequence present for at least 9 of the 10 non-elephant species. We tested the resulting 676,509 regions for acceleration in each elephant species relative to the 10 non-elephant species with phyloP using the following options: mode = ‘ACC’. We used the Q-Value method^81^ to adjust for multiple testing. Statistically significant ARs were defined with a false discovery rate threshold of 10%. We defined regions significantly accelerated in the Asian elephant, but not the African bush elephant as Asian elephant specific ARs and conversely defined African bush elephant specific ARs. Our previous studies of accelerated regions suggest no significant relationship between genome quality and number of ARs discovered^19^.

To define genes differentially expressed between Asian and African elephants we took advantage of the closeness between the two species. The Asian elephant is more closely related to the African elephant than humans are to chimpanzees (0.01186 substitutions / 100 bp vs 0.01277 substitutions / 100 bp based on fourfold degenerate sites from our WGA). For the purpose of defining differentially expressed genes, chimpanzee RNA-Seq reads have been aligned to human transcriptome indices^82^. We aligned African bush elephant PBMC reads (four technical replicates) from a previous study^19^ and publicly available Asian elephant PBMC data from a single individual^28^ (one replicate) to an African elephant (loxAfr3) transcriptome index with the STAR aligner^83^. After counting reads for each elephant gene with featureCounts^84^, we normalized counts with the TMM method and defined significant DE genes with edgeR^85^ (FDR < 0.01) correcting for multiple testing with the Benjamini-Hochberg method. The DE gene list was minimally affected by modest FDR cutoff changes. We note differences in the cell preps, RNA purification methods and sex of the Asian and African elephants as potential confounding factors in defining DE genes. The African elephant PBMC RNA was purified with a Ribo-Zero depletion step while the Asian elephant RNA was purified by Poly-A selection. A study comparing the two RNA purification methods shows a high gene expression correlation (0.931) between the two methods and detects 410 genes as differentially expressed when contrasting these purification methods^86^.

Potential regulatory regions for elephant DE genes were defined with custom R scripts implementing logic detailed by McLean et al. (2010)^87^ based on transcription start site (TSS) locations obtained for protein coding genes with the R package biomaRt^88^ for the African bush elephant genome (loxAfr3) with basal distances of 5 kb upstream and 1 kb downstream an extension distance of 100 kb. We chose this extension distance because the majority of conserved enhancers are located within 100 kb of a TSS^89^. We used the R package LOLA^90^ to test for enrichment of ARs relative to CRs in the potential regulatory regions of DE genes in the loxAfr3 genome. Biological processes (BP) and associated elephant orthologs of human genes were obtained with biomaRt. The resulting p-values were q-value corrected for multiple testing^81^. We used the same potential regulatory regions and LOLA to test for GO enrichments.

We compared elephant AR set GO enrichments to GO enrichments from previously published AR sets for 5 mammalian species (13-lined ground squirrel, naked mole rat, orca, bottlenose dolphin, and little brown bat)^19^. These AR sets were lifted over from hg19 coordinates to loxAfr3 coordinates. Numbers of ARs and background CRs overlapping potential regulatory regions of genes included in and excluded from each GO term were calculated with LOLA. We used generalized linear models with binomial distributions to compare elephant AR enrichments in each GO term to AR enrichments for the 5 other mammals. We contrasted models without and with an interaction term distinguishing the elephant AR set from the others. The two models are

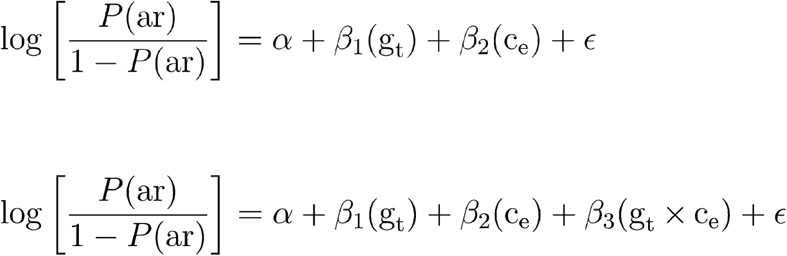

where *g_t_* is a binary value {0,1} indicating gene regions excluded from or included in a given GO term set; *c_e_* is a binary value {0,1} indicating non-elephant or elephant accelerated regions study; *ar* is the number of ARs in a given category. For each GO term with significant AR enrichments for at least one of the three elephant AR sets in the earlier analysis, we determined the significance of the enrichment in each elephant AR set relative to the other mammal AR sets by comparing the two models by likelihood ratio test. The likelihood ratio test p-values are reported in the Supplementary Data.

### Whole genome sequence analysis of living elephants

We obtained ~15–40x whole-genome sequencing data from multiple individuals from across the modern range of living elephants from public databases^12,16,28,91^, and the WGS libraries for “Hi-Dari” and “Icky” as well as a third African elephant named “Christie” (Supplementary Table 7). We also obtained sequence data from a straight-tusked elephant^25^ and two woolly mammoths^91^. Sequences were quality checked using FastQC and trimmed to remove nucleotide biases and adapter sequences with Trimmomatic where necessary. Reads from each individual were mapped to the *L. africana* genome (loxAfr3.0, Ensembl version) using bwa-mem v077^92^. Alignments were filtered to include only mapped reads and sorted by position using Samtools v0.0.19^93^, and we removed PCR duplicates using MarkDuplicates in picard v1.125^94^. Single-end reads from the ancient samples were mapped to loxAfr3.0 with bwa-aln following Palkolpoulou et al. (2018).

We estimated the number of *TP53* paralogs present in the genome of each elephant by calculating the average read depth of annotated sites in Ensembl *TP53* exons and whole genes with Samtools, dividing the total average genome coverage, multiplied by the number of TP53 annotations (n=12). We called variants in the living elephant species (n=13) by incorporating the .bam files using freebayes v1.3.1-12^36^, with extensive filtering to avoid false positives as follows with vcffilter from vcflib (https://github.com/vcflib/vcflib, last accessed July 2019): Phred-scale probability that a REF/ALT polymorphism exists at a given site (QUAL) > 20, the additional contribution of each observation should be 10 log units (QUAL/AO>10), read depth (DP>5), reads must exist on both strands (SAF>0 & SAR>0), and at least two reads must be balanced to each side of the site (RPR>1 & RPL>1). We then removed indels from the .vcf file and filtered to only include biallelic SNPs that were genotyped in every individual using VCFtools v0.1.17^95^ (-- remove-indels --min-alleles 2 --max-alleles 2 --max-missing-count 0) and bcftools v1.9^96^ (-v snps -m 1). We annotated the biallelic SNPs using SnpEff v4.3^97^ based on loxAfr3 (Ensembl), and calculated diversity statistics including per-individual heterozygosity, nucleotide diversity and Tajima’s D in 10kb windows with VCFtools, and the fixation index *F_ST_* with PopGenome v2.7.1^98^. We estimated RoH with PLINK v1.9^99^ using the following parameter settings: *-- homozyg-window-snp* 100 *--homozyg-window-missing* 15 *--homozyg-window-het* 5 *--homozyg-window-threshold* 0.05 *--homozyg-snp* 25 *--homozyg-kb* 100 *--homozyg-density* 50 *--homozyg-gap* 1000 *--homozyg-het* 750 *--allow-extra-chr*. For each elephant, the inbreeding coefficient (*F_ROH_*)^100^ was estimated using the total length of RoH≥500 Kb.

### Selective sweep analysis

To detect loci that have been putatively subjected to positive selection within each living elephant species, we used SweeD v3.3.1. SweeD scans the genome for selective sweeps by calculating the composite likelihood ratio (CLR) test from the folded site frequency spectrum in 1 kb grids across each scaffold. We used the folded site frequency spectrum because we lacked a suitable closely related outgroup with genomic resources that would enable us to establish ancestral alleles. For this analysis, we studied each species individually. Following Nielsen et al. (2005), we established statistical thresholds for this outlier analysis. First, we generated a null model by simulating 1,000 data sets with 100,000 SNPs under neutral demographic models. As our population genomics statistics results were highly concordant with Palkopoulou et al. (2018), we constructed demographic models based on their Pairwise Sequential Markovian Coalescent trajectories^101^ for each species, which we implemented with ms (October 2007 release)^102^ (Supplementary Fig. 9). Then, we categorized regions as outlier regions in the observed SNP data if their CLR was greater than the 99.99^th^ percentile of the distribution of the highest CLRs from the simulated SNP data. For the neutral simulations, we assumed a per-year mutation rate of 0.406e-09 and a generation time of 31 years, following Palkopoulou et al. (2018). We then calculated the CLR with the simulated neutral SNP datasets. SweeD output files were changed to BED format using namedCapture^103^ and data.table^104^ R packages, and we used bedtools intersect^105^ to collect elephant gene annotations (loxAfr3.0, Ensembl) overlapping putative selective sweeps.

Genomic scans for selection may be complicated by several factors that can increase false positive rates, and false negative rates potentially stem from variable mutation and recombination landscapes^38,106^. We established statistical thresholds using null demographic models. However, the estimated split times within living elephant species differ widely, ranging from 609,000 to 463,000 years before present for forest elephants^37^, to as recent as 38,000 to 30,000 years before present for bush elephants^91,107^. Estimated split times between the sampled Asian elephants are highly variable, ranging from 190,000 to 24,000 years before present^25^, indicating continental-wide population structure not accounted for here^108,109^. Still, Palkopoulou et al. (2018) found little evidence for gene flow between the three modern species of elephant, which supports our choice of analyzing them separately for selective sweeps. We focused on shared outlier regions, which show consistent evidence of being targeted by positive selection across all three elephant species.

Genes overlapping outlier regions of putative selective sweeps were functionally annotated by testing for GO enrichment of terms for biological processes^110^ in the outlier gene list, using default parameters and the Benjamini-Hochberg correction for multiple testing with an adjusted p-value < 0.05. We used REVIGO^111^ to semantically cluster and visualize the most significant GO terms according to their adjusted p-values using default parameters. We also created annotation clusters from the outlier genes using DAVID v6.8^112^ and constructed protein interaction networks with STRING v11.0^113^. Finally, we tested for enriched mouse phenotypes using ModPhea^114^.

## Data Availability

Short-read sequence data generated for this study has been shared under NCBI Bioproject PRJNA622303, and the genome assembly for Icky the Asian elephant is available on NCBI (GCA_014332765.1). Other datasets including the updated African elephant genome assembly, annotation files for Asian elephant, multiple genome alignments, *TP53* alignments and phylogeny, .vcf files, and selective sweep results have been deposited to Zenodo (https://zenodo.org/record/4033444#.X5cISFNKhGp).

## Acknowledgements

We acknowledge Leigh Duke for data coordination and Trent Fowler and Rosann Robinson for assistance with sample collection. We acknowledge the collections and veterinary staff at the San Diego Zoo Safari Park and Utah’s Hogle Zoo for sample collection. We acknowledge the following institutions for sharing data and/or resources: Buffalo Zoo, Dallas Zoo, El Paso Zoo, Fort Worth Zoo, Gladys Porter Zoo, Greenville Zoo, Jacksonville Zoo and Gardens, Louisville Zoological Garden, Oakland Zoo, Oklahoma City Zoo and Botanical Garden, Omaha’s Henry Doorly Zoo and Aquarium, The Phoenix Zoo, Point Defiance Zoo and Aquarium, San Antonio Zoological Society, Santa Barbara Zoological Gardens, Sedgwick County Zoo, Seneca Park Zoo, Toledo Zoological Gardens, Utah’s Hogle Zoo, Woodland Park Zoo, Zoo Atlanta, Zoo Miami and three other anonymous zoos. We acknowledge Huntsman Cancer Institute’s High-Throughput Genomics Core and the Monsoon computing cluster at Northern Arizona University (https://nau.edu/high-performance-computing/). Research reported in this publication was supported by the National Cancer Institute of the National Institutes of Health under Award Number U54CA217376. The content is solely the responsibility of the authors and does not necessarily represent the official views of the National Institutes of Health.

