## Supplementary Materials for "Elephant Genomes Elucidate Disease Defenses and Other Traits"

### Supplementary Data 1: ElephantGO_BP_EnrichmentsWithSpeciesComparisonQV.xlsx

We tested the potential cis-regulatory regions of Biological Process gene ontology genes for enrichments of elephant ARs relative to CRs. The column ‘GO_ID’ gives the gene ontology term enriched with ARs for at least one of the three elephant AR sets. The column ‘GO Term Description’ describes that GO term. ‘‘Elephant’ indicates the elephant AR set for a given row. ‘ElephantOddsRatio’ gives the odds ratio calculated with the R package LOLA for the overlap of elephant ARs relative to CRs with the potential regulatory regions of the GO genes.

‘ELephantLOLA_PValue’ indicates the p-value for the elephant AR set and GO term overlap. We also present a Q-value corrected false discovery rate in the column ‘ELephantLOLA_QValue’. LRT_PValue indicates the P-Value for the likelihood ratio test we performed contrasting the elephant AR–GO term overlap with the GO term overlaps for five other mammalian AR sets. We also present a Q-value corrected false discovery rate in the column ‘LRT_QValue’. The remaining columns give the LOLA calculated odds ratios for the GO term overlaps for AR sets from the 5 other mammals.

### Supplementary Information

#### **New elephant reference genomes**

We sequenced an estimated 94.4x coverage of the Asian elephant genome using multiple Illumina sequence libraries (Supplementary Table 1). These were assembled into a 3,126,981,281 bp draft reference genome organized on 6,954 scaffolds with a scaffold N50 of 2.78 Mb (Supplementary Table 2) and 91.5% of 4,104 mammalian single-copy orthologs in their complete form. We annotated 23,277 protein coding genes on 2,541 scaffolds, and 53% of the assembly was comprised of repeats (Supplementary Table 4).

We also used Hi-C sequencing libraries to improve the African bush elephant genome assembly, increasing the length of the longest scaffold from 225 Mb to 240 Mb (Supplementary Table 5). A total of 528 previously unjoined scaffolds were linked and nine scaffolds/chromosomes were broken (Supplementary Fig. 1). We suggest future improvements to these assemblies such as high-resolution genomic profiling with long read and/or optical mapping technologies to further resolve structural differences between elephant genomes.

### Supplementary Figure 1.Tuberculosis (TB) rates in Asian and African elephants. In a study of captive elephants, Asian elephants tested positive for TB in significantly higher numbers than did African elephants^1^. Shown is the results from the Chi-squared test (P=0.0006843).

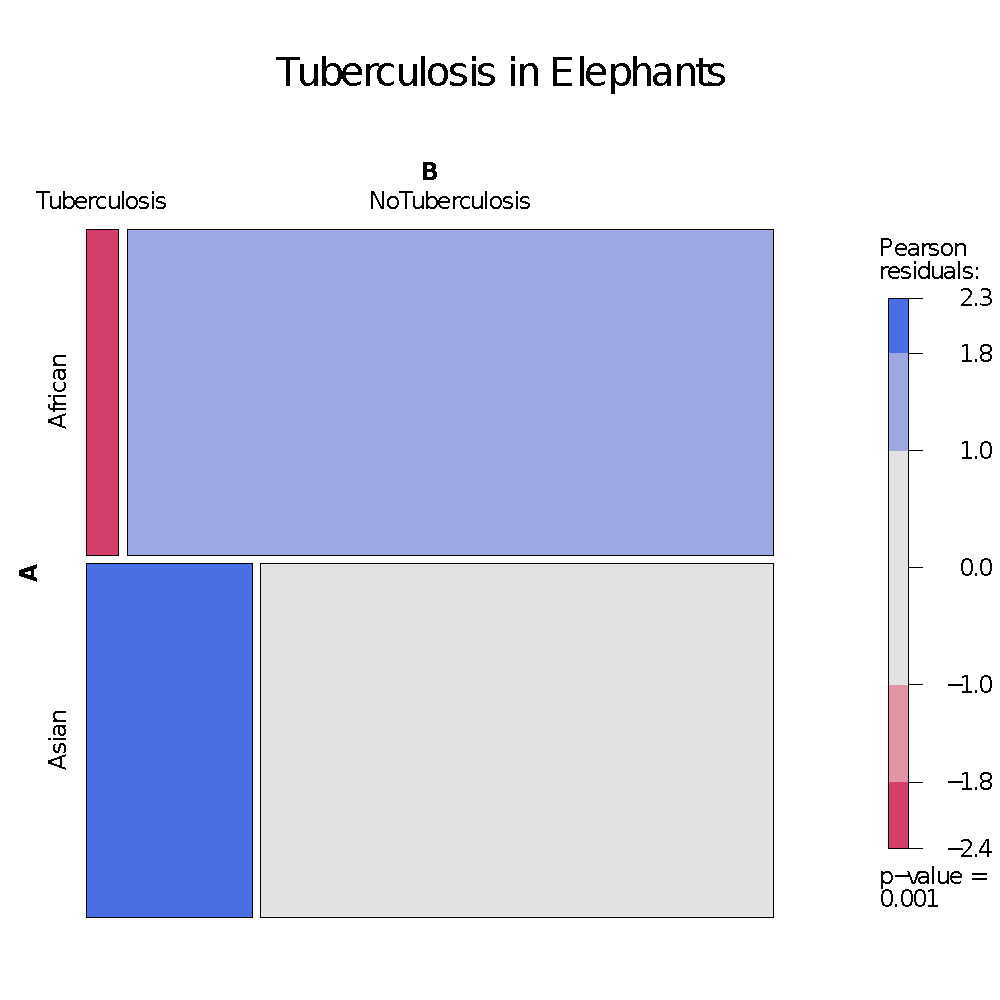

### Supplementary Figure 2. Synteny analyses between the Hi-C and loxAfr4.0 African bush elephant assemblies. (a) Jupiter plot showing correspondence between assemblies considering the total length of both reference and query assemblies. Plotting tool from https://github.com/JustinChu/JupiterPlot. (b) Dot plot of the percent identity and mapping positions between scaffolds in the two assemblies from minimap^2^.

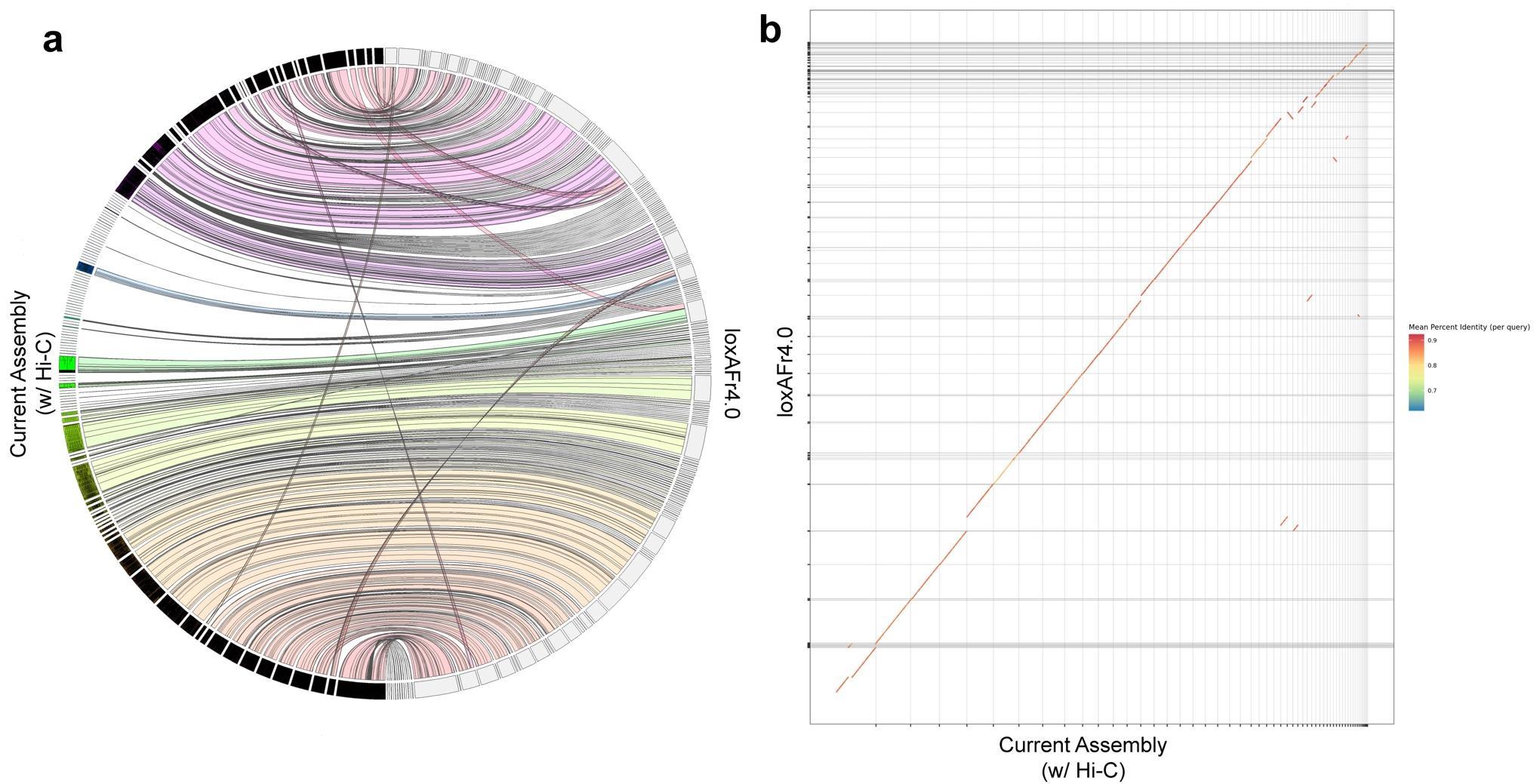

### Supplementary Figure 3. Species-specific accelerated regions (ARs) disproportionately overlap highly differentially expressed gene regulatory regions relative to the common ARs. This pattern is robust to a range of adjusted p-values. Odds ratio (a) and -log10(p-value) (b).

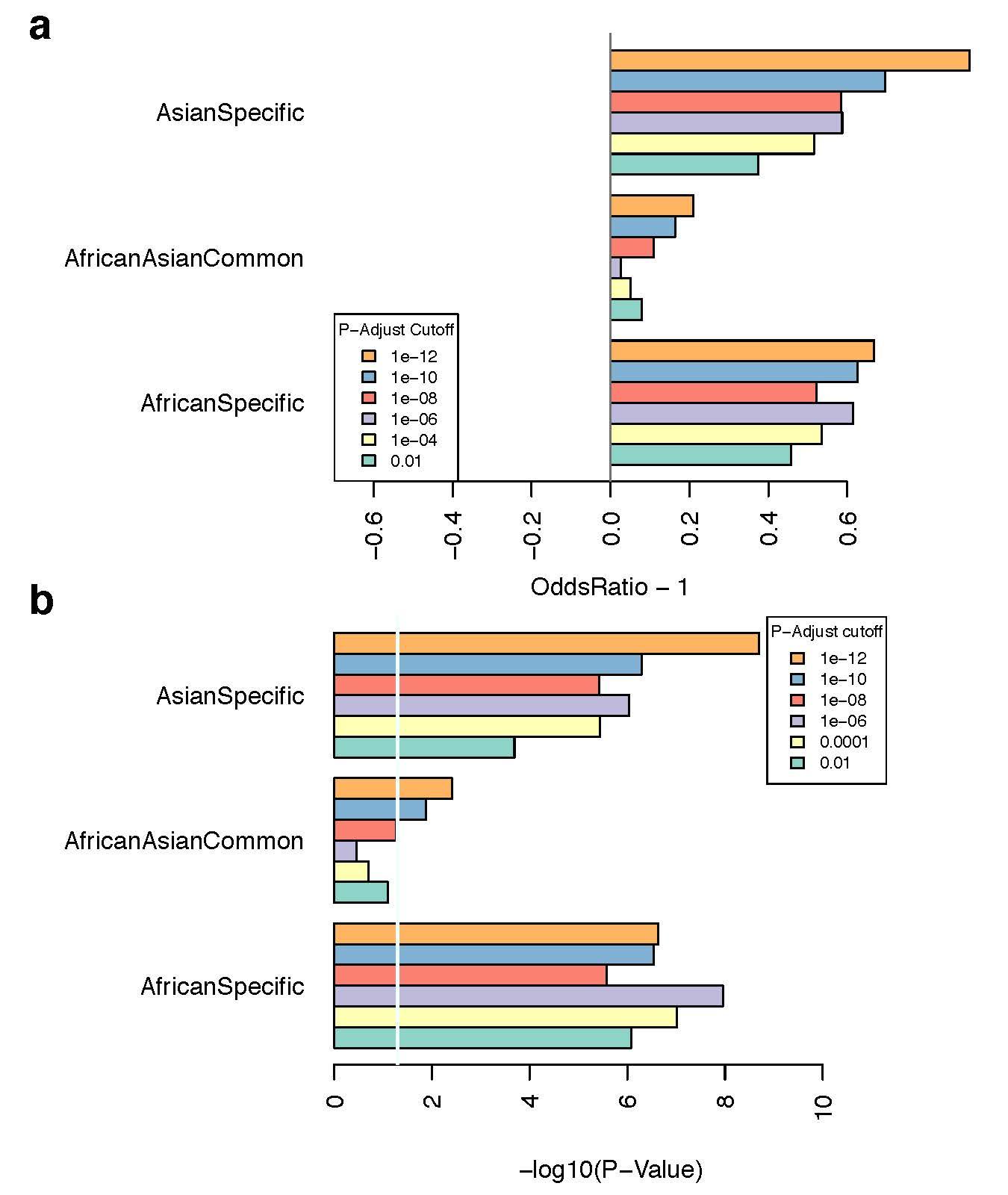

### Supplementary Figure 4. Semantic clustering of significantly enriched Gene Ontology terms for biological processes found near Asian elephant-specific accelerated regions. Rectangles represent clusters, and larger rectangles indicate semantically related clusters. Larger rectangle sizes reflect smaller corrected p-values from the GO term enrichment.

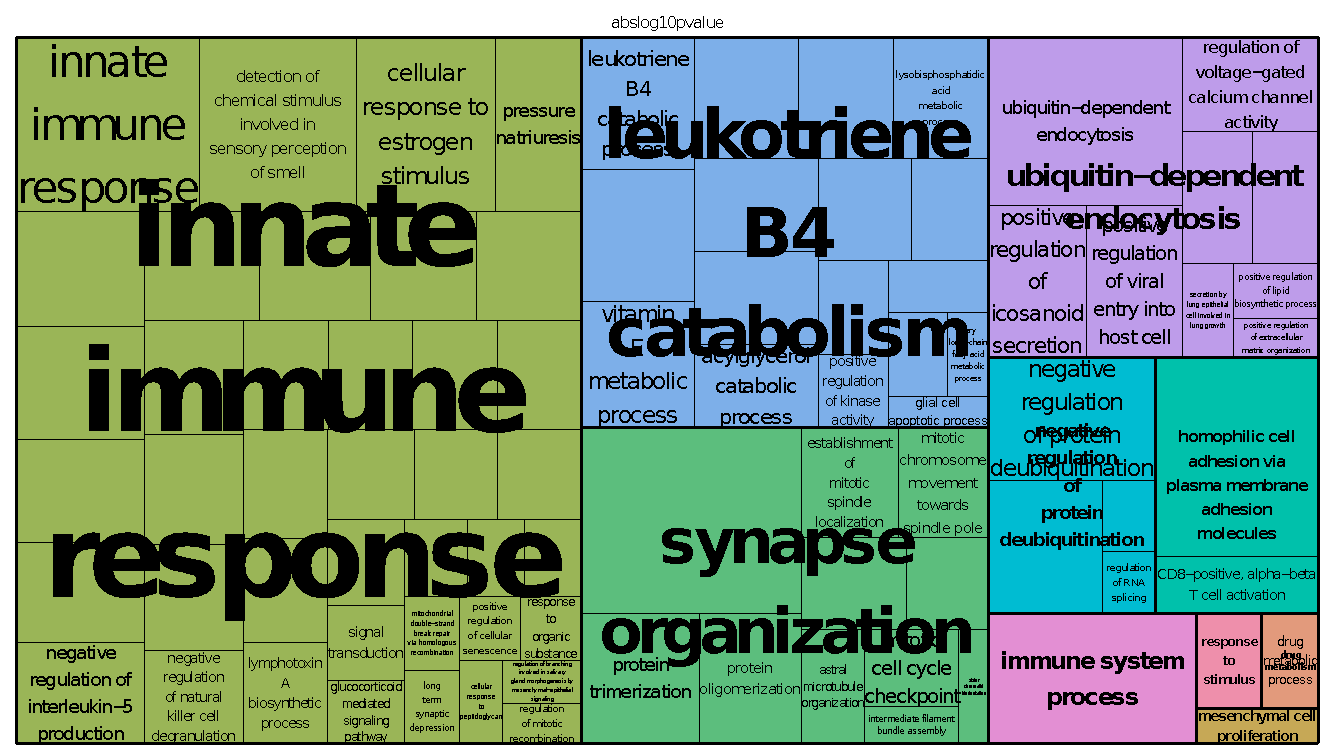

### Supplementary Figure 5. Semantic clustering of significantly enriched Gene Ontology terms for biological processes found near African bush elephant-specific accelerated regions. Rectangles represent clusters, and larger rectangles indicate semantically related clusters. Larger rectangle sizes reflect smaller corrected p-values from the GO term enrichment.

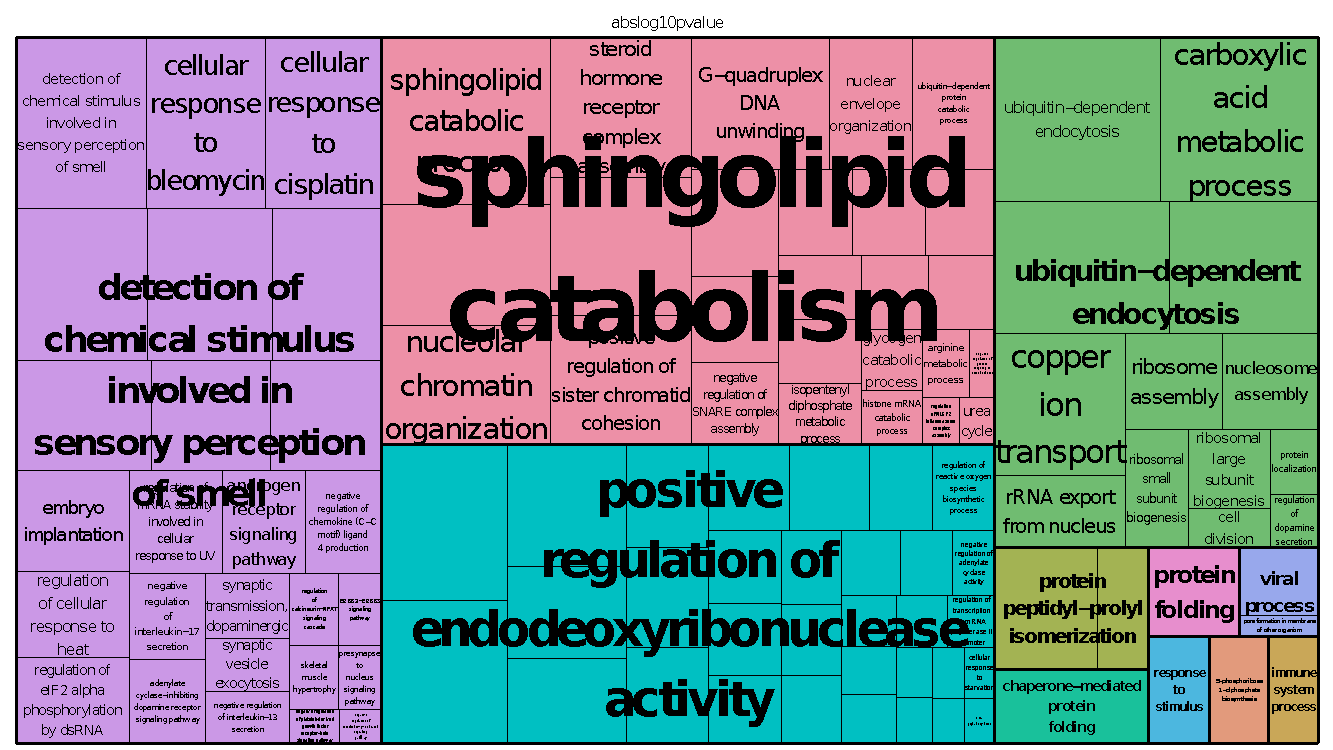

### Supplementary Figure 6. Semantic clustering of significantly enriched Gene Ontology terms for biological processes found near common elephant accelerated regions. Rectangles represent clusters, and larger rectangles indicate semantically related clusters. Larger rectangle sizes reflect smaller corrected p-values from the GO term enrichment.

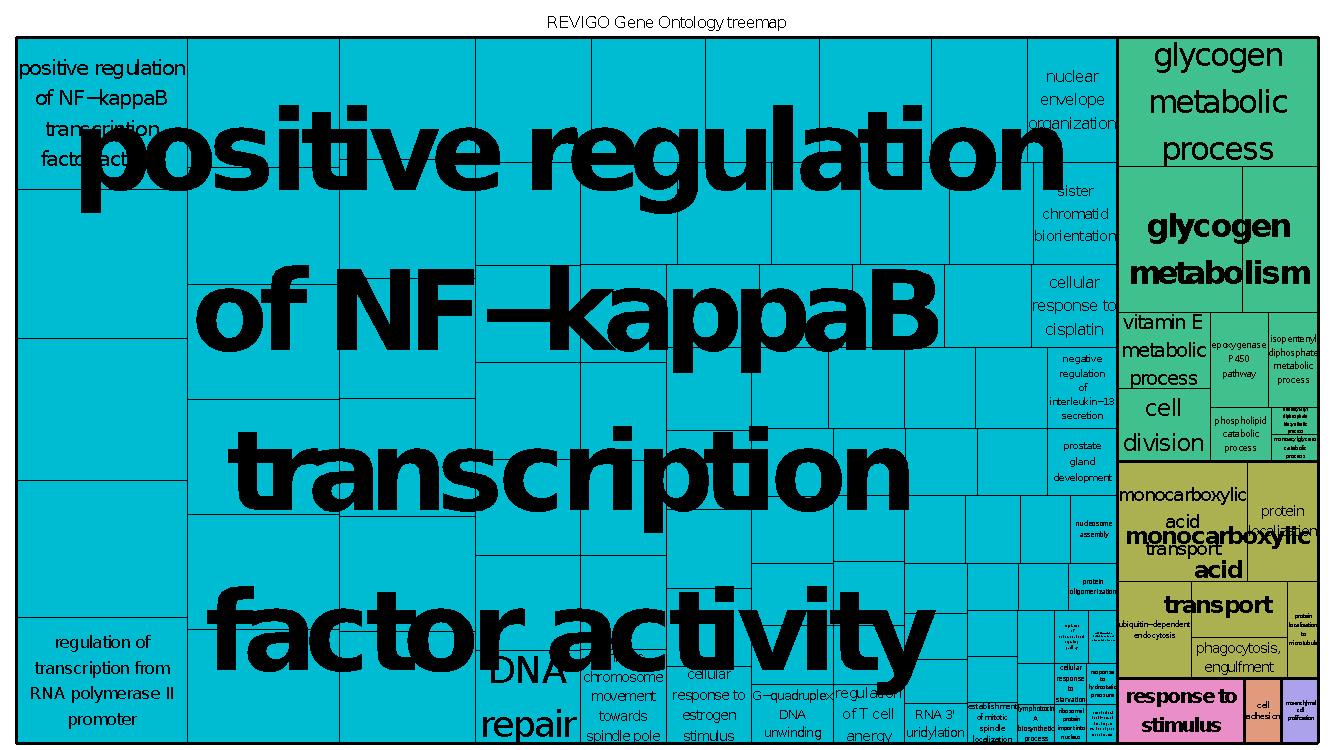

### Supplementary Figure 7. Evolution of TP53 in mammals. Phylogenetic reconstruction using BEAST2^3^ of TP53 homologs from 44 mammalian genome assemblies. Node labels indicate posterior probabilities. Node bars indicate 95% highest posterior density for node age. Scale axis in millions of years.

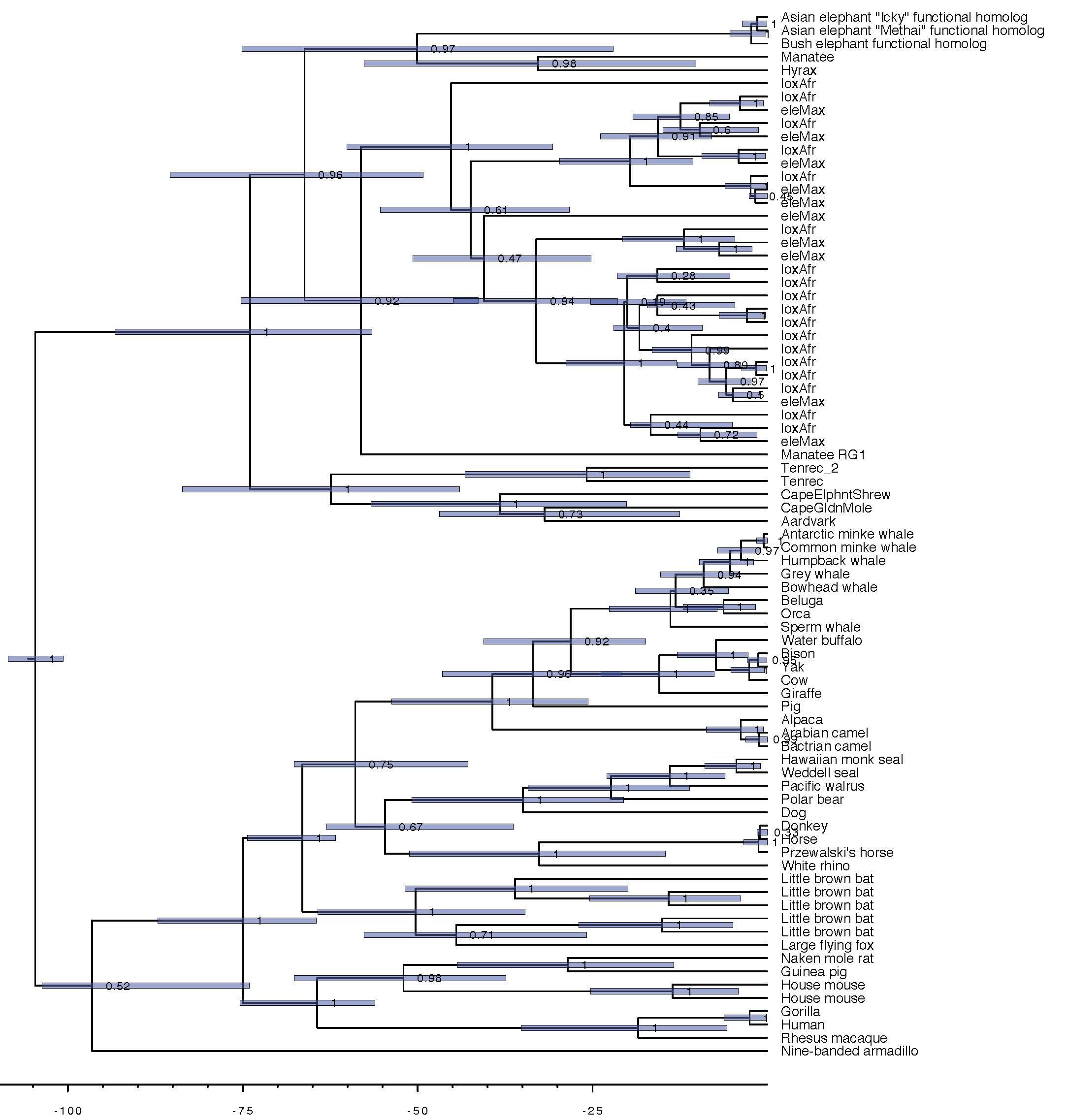

### Supplementary Figure 8. Runs of homozygosity (RoH) in elephant genomes. (a) Frequency distributions of RoH in Kb across 13 individual elephant genomes. (b) Inbreeding coefficients (*F_ROH_*) for 13 elephant genomes.

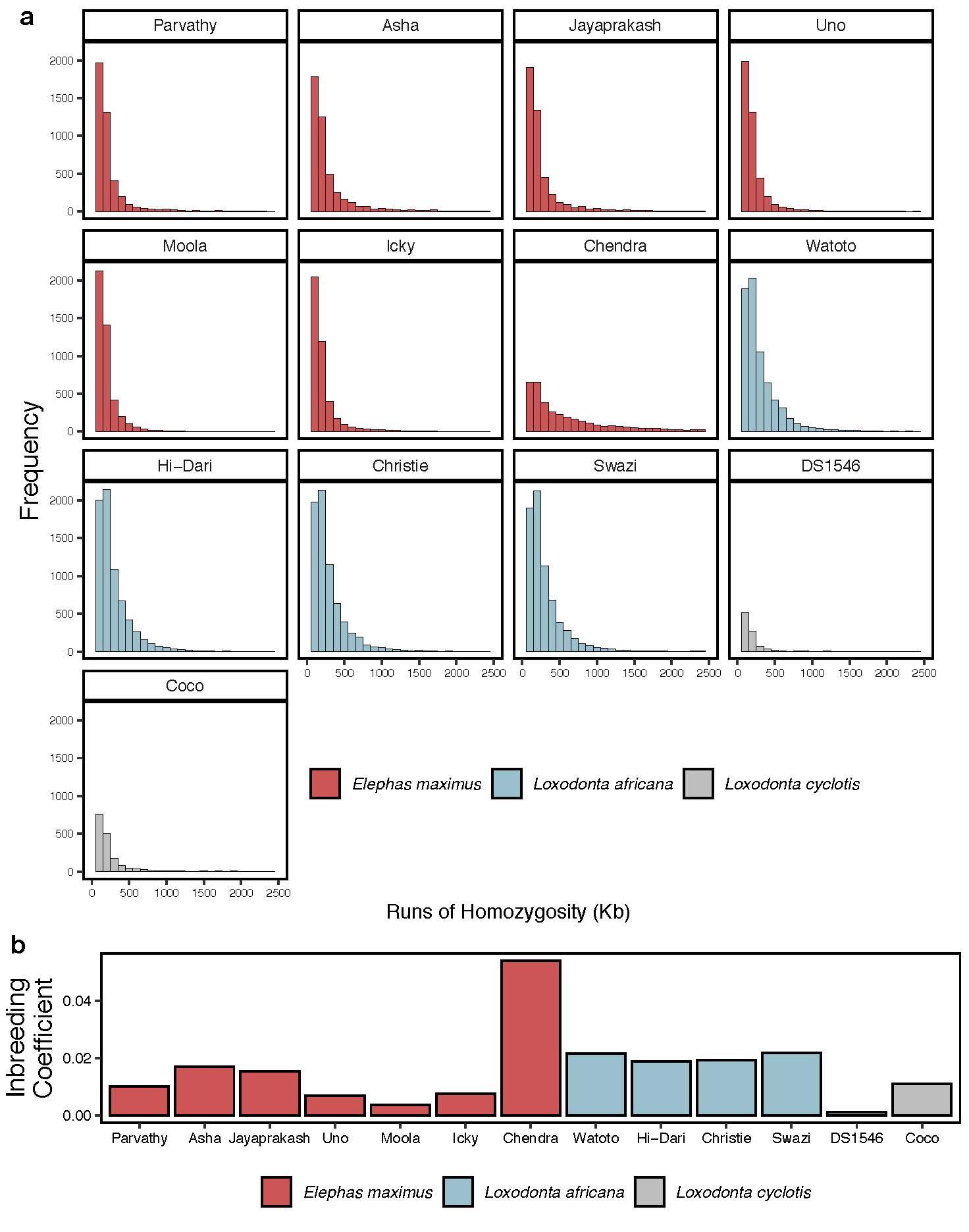

### Supplementary Figure 9. Demographic models for three elephant species, implemented in Hudson’s ms^4^. Width of grey bars represent population size changes over time. (a) Asian elephant (*Elephas maximus*): 14 100 -s 100000 -t 0.01 -G -1.24 -eN 1 17.5 -eG 2 0.14 -eG 4 -0.096 -eN 37.5 25. (b) African bush elephant (*Loxodonta Africana*): 8 100 -s 100000 -t 10 -eN 4.8 352 -eG 29 .07. (c) African forest elephant (*L. cyclotis*): 4 100 -s 100000 -I 2 2 2 -n 2 4.0 -en 0.16 1 4.0 -en 0.33 2 13.4 -en 1.6 2 4.0 -ej 1.6 1 2 -en 6.45 2 13.4 -en 33.0 2 100.0.

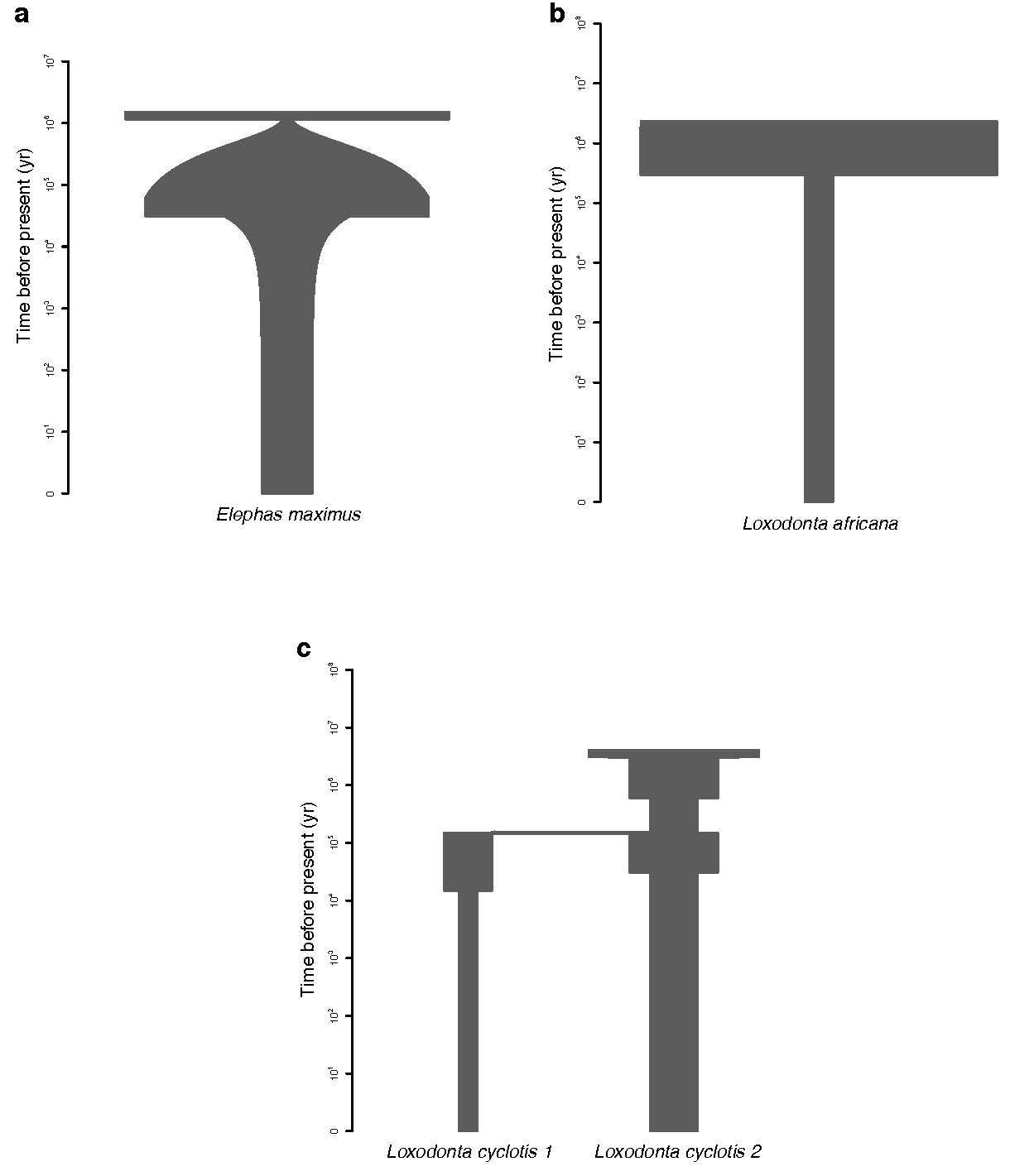

### Supplementary Table 1. Demographics of elephants with and without records of neoplasia

| **No record of neoplasia/cancer** | | | | |
| --- | --- | --- | --- | --- |
| **Species** | **Common Name** | **Sex** | **Age** | **Necropsy** |
| *Loxodonta africana* | AFRICAN ELEPHANT | male | 28 | no |
| *Loxodonta africana* | AFRICAN ELEPHANT | female | 43 | no |
| *Loxodonta africana* | AFRICAN ELEPHANT | female | 31 | no |
| *Loxodonta africana* | AFRICAN ELEPHANT | female | 33 | no |
| *Loxodonta africana* | AFRICAN ELEPHANT | female | 30 | no |
| *Loxodonta africana* | AFRICAN ELEPHANT | female | 49 | yes |
| *Loxodonta africana* | AFRICAN ELEPHANT | male | 3 | yes |
| *Loxodonta africana* | AFRICAN ELEPHANT | female | 43 | yes |
| *Loxodonta africana* | AFRICAN ELEPHANT | female | 36 | no |
| *Loxodonta africana* | AFRICAN ELEPHANT | female | 38 | no |
| *Loxodonta africana* | AFRICAN ELEPHANT | male | 26 | no |
| *Loxodonta africana* | AFRICAN ELEPHANT | female | 45 | no |
| *Loxodonta africana* | AFRICAN ELEPHANT | male | 7 | no |
| *Loxodonta africana* | AFRICAN ELEPHANT | female | 27 | yes |
| *Loxodonta africana* | AFRICAN ELEPHANT | male | 38 | no |
| *Loxodonta africana* | AFRICAN ELEPHANT | male | 40 | yes |
| *Loxodonta africana* | AFRICAN ELEPHANT | female | 20 | yes |
| *Loxodonta africana* | AFRICAN ELEPHANT | female | 45 | yes |
| *Loxodonta africana* | AFRICAN ELEPHANT | female | 47 | yes |
| *Loxodonta africana* | AFRICAN ELEPHANT | female | 39 | yes |
| *Loxodonta africana* | AFRICAN ELEPHANT | female | 45 | no |
| *Loxodonta africana* | AFRICAN ELEPHANT | female | 29 | no |
| *Loxodonta africana* | AFRICAN ELEPHANT | female | 26 | no |
| *Loxodonta africana* | AFRICAN ELEPHANT | female | 31 | no |
| *Loxodonta africana* | AFRICAN ELEPHANT | female | 3 | yes |
| *Loxodonta africana* | AFRICAN ELEPHANT | female | 20 | no |
| *Loxodonta africana* | AFRICAN ELEPHANT | female | 24 | no |
| *Loxodonta africana* | AFRICAN ELEPHANT | male | 32 | yes |
| *Loxodonta africana* | AFRICAN ELEPHANT | female | 55 | yes |
| *Loxodonta africana* | AFRICAN ELEPHANT | NA | NA | yes |
| *Loxodonta africana* | AFRICAN ELEPHANT | NA | NA | yes |
| *Loxodonta africana* | AFRICAN ELEPHANT | NA | NA | yes |
| *Loxodonta africana* | AFRICAN ELEPHANT | NA | NA | yes |
| *Elephas maximus* | ASIAN ELEPHANT | female | 28 | no |
| *Elephas maximus* | ASIAN ELEPHANT | female | 25 | NA |
| *Elephas maximus* | ASIAN ELEPHANT | female | 51 | no |
| *Elephas maximus* | ASIAN ELEPHANT | female | 5 | yes |
| *Elephas maximus* | ASIAN ELEPHANT | female | 37 | yes |
| *Elephas maximus* | ASIAN ELEPHANT | female | 35 | no |
| *Elephas maximus* | ASIAN ELEPHANT | female | 36 | yes |
| *Elephas maximus* | ASIAN ELEPHANT | female | 20 | yes |
| *Elephas maximus* | ASIAN ELEPHANT | female | 47 | no |
| *Elephas maximus* | ASIAN ELEPHANT | female | 45 | no |
| *Elephas maximus* | ASIAN ELEPHANT | female | 33 | NA |
| *Elephas maximus* | ASIAN ELEPHANT | female | 15 | no |
| *Elephas maximus* | ASIAN ELEPHANT | female | 28 | no |
| *Elephas maximus* | ASIAN ELEPHANT | male | 40 | yes |
| *Elephas maximus* | ASIAN ELEPHANT | female | 41 | yes |
| *Elephas maximus* | ASIAN ELEPHANT | female | 47 | no |
| *Elephas maximus* | ASIAN ELEPHANT | female | 50 | yes |
| *Elephas maximus* | ASIAN ELEPHANT | male | 31 | no |
| *Elephas maximus* | ASIAN ELEPHANT | female | 49 | no |
| *Elephas maximus* | ASIAN ELEPHANT | NA | NA | yes |
| *Elephas maximus* | ASIAN ELEPHANT | NA | NA | yes |
| *Elephas maximus* | ASIAN ELEPHANT | NA | NA | yes |
| *Elephas maximus* | ASIAN ELEPHANT | NA | NA | yes |
| *Elephas maximus* | ASIAN ELEPHANT | NA | NA | yes |
| **Record of neoplasia/cancer** | | | | |
| **Species** | **Common Name** | **Sex** | **Age** | **Necropsy** |
| *Loxodonta africana* | AFRICAN ELEPHANT | female | 28 | no |
| *Loxodonta africana* | AFRICAN ELEPHANT | NA | NA | yes |
| *Elephas maximus* | ASIAN ELEPHANT | female | 45 | NA |
| *Elephas maximus* | ASIAN ELEPHANT | female | 50 | yes |
| *Elephas maximus* | ASIAN ELEPHANT | female | 30, 40 | no |
| *Elephas maximus* | ASIAN ELEPHANT | female | 39 | NA |
| *Elephas maximus* | ASIAN ELEPHANT | female | 39 | no |
| *Elephas maximus* | ASIAN ELEPHANT | male | 35 | no |
| *Elephas maximus* | ASIAN ELEPHANT | female | 50 | no |
| *Elephas maximus* | ASIAN ELEPHANT | female | 36 | no |
| *Elephas maximus* | ASIAN ELEPHANT | female | 50 | no |
| *Elephas maximus* | ASIAN ELEPHANT | female | 59 | yes |
| *Elephas maximus* | ASIAN ELEPHANT | NA | NA | yes |
| *Elephas maximus* | ASIAN ELEPHANT | NA | NA | yes |
| *Elephas maximus* | ASIAN ELEPHANT | NA | NA | yes |
| *Elephas maximus* | ASIAN ELEPHANT | NA | NA | yes |
| *Elephas maximus* | ASIAN ELEPHANT | NA | NA | yes |
| *Elephas maximus* | ASIAN ELEPHANT | NA | NA | yes |
| *Elephas maximus* | ASIAN ELEPHANT | NA | NA | yes |

### Supplementary Table 2. Genomic Sequence Data Obtained for the Asian Elephant.

| **Library insert size and type** | **Read lengths** | **Coverage** |
| --- | --- | --- |
| 200 bp paired-end | 2x125 bp | 34x |
| 3 kb mate-paired | 2x100 bp | 15x |
| 5 kb mate-paired | 2x100 bp | 13.8x |
| 8 kb mate-paired | 2x100 bp | 14.7x |
| 10 kb mate-paired | 2x100 bp | 15.2x |
| Total coverage: | | 94.4x |

### Supplementary Table 3. Summary Statistics for the Asian Elephant Genome Assembly.

| **Feature** | **Contigs** | **Scaffolds** |
| --- | --- | --- |
| Assembly length | 2.98Gb | 3.13 Gb |
| Longest | 731 kb | 14.6 Mb |
| Number | 90,662 | 6,954 |
| N50 | 79.8 kb | 2.77 Mb |
| L50 | 10,736 | 336 |
| Percent genome in gaps | 0.09 | 4.88 |
| BUSCO results | C: 91.5% [D:0.4%], F:5.7%, M:2.8%, n:4,104 | |

BUSCO: Benchmarking Universal Single Copy Orthologs; C: complete; D: duplicated; F: fragmented; M: missing

### Supplementary Table 4. Interspersed Repeat Content of the Asian Elephant Genome Assembly, Estimated with a Library of Known Mammalian Repeats (RepBase) and De Novo Repeat Identification (RepeatModeler).

| **Repeat Type** | **RepBase** | | **RepeatModeler** | |
| --- | --- | --- | --- | --- |
|  | **Length (bp)** | **% Genome (51.52% total)** | **Length (bp)** | **% Genome (46.11 total)** |
| SINEs | 342,400,474 | 10.94 | 55,270,845 | 1.77 |
| LINEs | 872,332,896 | 27.88 | 1,031,262,873 | 32.96 |
| LTR | 241,033,286 | 7.70 | 195,360,272 | 6.24 |
| DNA transposons | 87,641,059 | 2.80 | 62,073,944 | 1.98 |
| Unclassified | 6,190 | 0.04 | 98,648,742 | 3.15 |

### Supplementary Table 5. Comparison of Statistics for the African Savannah Elephant Genome Assemblies.

|  | **loxAfr3** | | **loxAfr4** | | **Current study** | |
| --- | --- | --- | --- | --- | --- | --- |
| **Feature** | **Contigs** | **Scaffolds** | **Contigs** | **Scaffolds** | **Contigs** | **Scaffolds** |
| Assembly length | 3.1 Gb | 3.2 Gb | 3.1 Gb | 3.3 Gb | 3. 1 Gb | 3.3 Gb |
| Longest | 567 kb | 129 Mb | 567 kb | 225 Mb | 567 kb | 240 Mb |
| Number | 95,867 | 2,353 | 95,891 | 2,303 | 95,889 | 1,784 |
| N50 | 69 kb | 46 Mb | 69 kb | 94 Mb | 69 kb | 89 Mb |
| L50 | 13,607 | 21 | 13,607 | 11 | 13,607 | 11 |
| Percent genome in gaps | 0 | 2.45 | 0 | 4.68 | 0 | 4.69 |

### Supplementary Table 6. Genome assemblies queried for TP53 homologs.

| **Species Name** | **Assembly ID** |
| --- | --- |
| *Vicugna pacos*  Alpaca | Vi_pacos_V1.0 |
| *Bison bison*  American Bison | Bison_UMD1.0 |
| *Balaenoptera bonaerensis*  Antarctic Minke Whale | ASM97880v1 |
| *Camelus dromedarius*  Arabian Camel | PRJNA234474_Ca_dromedarius_V1.0 |
| *Dasypus novemcinctus*  Nine-Banded Armadillo | Dasnov3.0 |
| *Camelus bactrianus*  Bactrian Camel | Ca_bactrianus_MBC_1.0 |
| *Delphinapterus leucas*  Beluga Whale | ASM228892v3 |
| Balaena mysticetus  Bowhead Whale | http://www.bowhead-whale.org/downloads/ |
| *Bos taurus*  Cattle | ARS-UCD1.2 |
| *Canis lupus familiaris*  Dog | CanFam3.1 |
| *Equus asinus asinus*  Donkey | ASM30337v1 |
| *Loxodonta africana*  African Savanna Elephant | Loxafr3.0 |
| *Giraffa tippelskirchi*  Giraffe | ASM165123v1 |
| *Gorilla gorilla gorilla*  Western Lowland Gorilla | GorGor4 |
| *Eschrichtius robustus*  Grey Whale | ASM218922v1 |
| *Cavia porcellus*  Domestic Guinea Pig | Cavpor3.0 |
| *Monachus schauinslandi*  Hawaiian Monk Seal | ASM220157v1 |
| *Equus caballus*  Horse | EquCab3.0 |
| *Homo sapiens*  Human | GRCh38.p13 |
| *Megaptera novaeangliae*  Humpback Whale | megNov1 |
| *Pteropus vampyrus*  Large Flying Fox | Pvam_2.0 |
| *Myotis lucifugus*  Little Brown Bat | Myoluc2.0 |
| *Trichechus manatus latirostris*  Florida Manatee | TriManLat1.0 |
| *Balaenoptera acutorostrata*  Minke Whale | BalAcu1.0 |
| *Mus musculus*  House Mouse | GRCm38.p6 |
| *Heterocephalus glaber*  Naked Mole-Rat | HetGla_female_1.0 |
| *Orcinus orca*  Killer Whale | Oorc_1.1 |
| *Sus scrofa*  Pig | Sscrofa11.1 |
| *Ursus maritimus*  Polar Bear | UrsMar_1.0 |
| *Equus przewalskii*  Przewalski’s Horse | Burgud |
| *Macaca mulatta*  Rhesus Monkey | rheMacS_1.0 |
| *Procavia capensis*  Cape Rock Hyrax | Pcap_2.0 |
| *Physeter catodon*  Sperm Whale | ASM283717v2 |
| *Odobenus rosmarus*  Pacific Walrus | Oros_1.0 |
| *Bubalus bubalis*  Water Buffalo | Bubbub1.0 |
| *Leptonychotes weddellii*  Weddell Seal | LepWed1.0 |
| *Ceratotherium simum*  Southern White Rhinoceros | CerSimSim1.0 |
| *Camelus ferus*  Wild Bactrian Camel | CB1 |
| *Bos mutus*  Wild Yak | BosGru_v2.0 |
| *Elephas maximus*  Asian elephant “Methai” | https://www.dnazoo.org/assemblies/Elephas_maximus |
| *Elephas maximus*  Asian elephant “Icky” | Current study |

### Supplementary Table 7. Summary of Elephant Whole-genome Shotgun Resequencing Data Utilized in This Study, Mapped to the African Bush Elephant Reference Assembly (loxAfr3.0).

| **Species** | **Name** | **Geographic Origin** | **Source** | **# Mapped Reads** | **Prop. Reads Properly Paired** | **Peak Read Depth** |
| --- | --- | --- | --- | --- | --- | --- |
| *Loxodonta africana* | Watoto | Kenya | ERR2260496 | 874,537,386 | 0.99 | 26X |
|  | Swazi | South Africa | ERR2260497 | 1,014,067,450 | 0.93 | 30X |
|  | HI-Dari | Kenya | SRR11869865,SRR11869866(Current study) | 1,072,817,612 | 0.97 | 38X |
|  | Christie | Zimbabwe | SRR12799664,  SRR12799663  (Current study, Abegglen et al. 2015) | 1,031,044,341 | 0.98 | 36X |
| *Loxodonta cyclotis* | DS1546 | Central African Republic | ERR2260495 | 852,948,500 | 0.96 | 24X |
|  | Coco | Sierra Leone | ERR2260500 | 981,145,080 | 0.99 | 30X |
| *Elephas maximus* | Moola | Myanmar | ERR2260498 | 1,188,021,033 | 0.9 | 36X |
|  | Chendra | Borneo | ERR2260499 | 981,228,188 | 0.99 | 30X |
|  | Icky | Myanmar | SRR11577048  (Current study) | 898,020,572 | 0.97 | 32X |
|  | Parvathy | India | SRR2008170 | 872,535,345 | 0.96 | 26X |
|  | Asha | India | SRR2009586 | 977,136,495 | 0.94 | 29X |
|  | Uno | Assam, India | SRR2012205, SRR2012206, SRR2012207 | 912,606,191 | 0.96 | 26X |
|  | Jayaprakash | Karnataka, India | SRR2912975 | 475,023,505 | 0.94 | 13X |
| *Palaeoloxodon antiquus* | NA | Germany | ERR2260504 | 916,662,984 | NA | 7X |
| *Mammuthus primigenius* | NA | Oimyakon, Russia | ERR852028 | 617,446,606 | NA | 10X |
|  | NA | Wrangel Island, Russia | ERR855944 | 760,223,385 | NA | 16X |

### Supplementary Table 8. Estimates of TP53 Copy Numbers in the Genomes of Living and Extinct (†) Elephant Species Based on Whole Genome Shotgun Data Mapping.

| **Species** | **Exons Only** | | | **Whole Gene** | | |
| --- | --- | --- | --- | --- | --- | --- |
|  | **range** | **mean** | **stdev** | **range** | **mean** | **stdev** |
| *Loxodonta africana* | 18.0–22.4 | 20.1 | 1.80 | 16.4–19.6 | 18.0 | 1.32 |
| *Loxodonta cyclotis* | 24.2–25.2 | 24.7 | 0.65 | 21.1–22.3 | 21.7 | 0.92 |
| *Elephas maximus* | 10.8–36.8 | 21.9, 21.1* | 8.01, 3.07* | 10.3–32.4 | 19.6, 19.0* | 6.81, 2.47* |
| *Palaeoloxodon antiquus^†^* | NA | 25.1 | NA | NA | 22.6 | NA |
| *Mammuthus primigenius^†^* | 21.0–28.0 | 24.5 | NA | 18.9–24.0 | 21.0 | NA |

*Descriptive statistics calculated after removing 2 outliers ≥1 standard deviation from the mean

### Supplementary Table 9. Genetic variation in TP53 paralogs estimated from three living elephant species compared to ancestral repeats.

| **Gene ID** | **No. sites** | **No. SNPs** | **Syn. SNPs** | **Nonsyn. SNPs** | ***Elephas maximus* (n=7)** | | ***Loxodonta Africana* (n=4)** | | ***Loxodonta cyclotis* (n=2)** | | ***F_ST_***  **(all SNPs)** | ***F_ST_***  **Syn.** | ***F_ST_* Nonsyn.** |
| --- | --- | --- | --- | --- | --- | --- | --- | --- | --- | --- | --- | --- | --- |
|  |  |  |  |  | **Seg. sites** | **Nuc. Diversity** | **Seg. sites** | **Nuc. Diversity** | **Seg. sites** | **Nuc. Diversity** |  |  |  |
| ENSLAFG00000026238 | 1126 | 4 | 2 | 2 | 0 | 0 | 1 | 0.00059 | 0 | 0 | 0.33 | 0 | 0.33 |
| ENSLAFG00000027820 | 1128 | 2 | 0 | 2 | 0 | 0 | 0 | 0 | 1 | 0.00089 | 0 | 0 | 0 |
| ENSLAFG00000030880 | 888 | 2 | 0 | 2 | 0 | 0 | 0 | 0 | 0 | 0 | 0 | 0 | 1 |
| ENSLAFG00000027669 | 1126 | 8 | 2 | 6 | 1 | 0.00025 | 0 | 0 | 0 | 0 | 0.88 | 0 | 1 |
| ENSLAFG00000027348 | 1130 | 6 | 1 | 4 | 3 | 0.00101 | 0 | 0 | 0 | 0 | 0.63 | 0 | 0.88 |
| ENSLAFG00000007483* | 5559 | 33 | 5 | 0 | 6 | 0.00039 | 0 | 0 | 0 | 0 | 0.91 | 1 | 0 |
| ENSLAFG00000030555 | 1127 | 5 | 2 | 3 | 1 | 0.00025 | 1 | 0.00044 | 0 | 0 | 0.65 | 0 | 0.83 |
| ENSLAFG00000027474 | 1140 | 17 | 8 | 8 | 3 | 0.00083 | 0 | 0 | 1 | 0 | 0.80 | 0.77 | 0.84 |
| ENSLAFG00000027365 | 1125 | 7 | 0 | 6 | 2 | 0.0008 | 0 | 0 | 0 | 0 | 0.75 | 0 | 0.75 |
| ENSLAFG00000032042 | 1126 | 1 | 0 | 1 | 0 | 0 | 0 | 0 | 1 | 0.0009 | 0.5 | 0 | 0.5 |
| ENSLAFG00000028692 | 1125 | 0 | 0 | 0 | 0 | 0 | 0 | 0 | 0 | 0 | 0 | 0 | 0 |
| ENSLAFG00000028299** | 1126 | 0 | 0 | 0 | 0 | 0 | 0 | 0 | 0 | 0 | 0 | 0 | 0 |
| Ancestral repeats | 18381865 | 232948 | NA | NA | 32422 | 0.0006 | 9334 | 0.00029 | 25766 | 0.00140 | 0 | NA | NA |

*Functional homolog

**”retrogene 9” from Abegglen et al. (2015)

### Supplementary Table 10. Number of annotated variants by effect and type in the Ensembl-annotated African bush elephant TP53 paralogs.

| **Gene ID** | **High Impact** | **Low Impact** | **Moderate Impact** | **Downstream Gene Effect** | **Introns** | **Missense** | **Splice Donor** | **Splice Region** | **Stop Codons Gained** | **Synonymous** | **Upstream Gene Effect** |
| --- | --- | --- | --- | --- | --- | --- | --- | --- | --- | --- | --- |
| ENSLAFG00000026238 | 1 | 2 | 1 | 3 | 0 | 1 | 0 | 0 | 1 | 2 | 10 |
| ENSLAFG00000027820 | 0 | 0 | 2 | 16 | 0 | 2 | 0 | 0 | 0 | 0 | 26 |
| ENSLAFG00000030880 | 0 | 0 | 2 | 12 | 0 | 2 | 0 | 0 | 0 | 0 | 20 |
| ENSLAFG00000027669 | 0 | 2 | 6 | 56 | 0 | 6 | 0 | 1 | 0 | 2 | 19 |
| ENSLAFG00000027348 | 0 | 2 | 4 | 43 | 1 | 4 | 0 | 1 | 0 | 1 | 30 |
| ENSLAFG00000007483* | 0 | 5 | 0 | 37 | 28 | 0 | 0 | 0 | 0 | 5 | 59 |
| ENSLAFG00000030555 | 1 | 2 | 2 | 35 | 0 | 2 | 0 | 0 | 1 | 2 | 14 |
| ENSLAFG00000027474 | 1 | 8 | 8 | 65 | 1 | 8 | 1 | 1 | 0 | 8 | 74 |
| ENSLAFG00000027365 | 0 | 1 | 6 | 37 | 1 | 6 | 0 | 2 | 0 | 0 | 28 |
| ENSLAFG00000032042 | 0 | 0 | 1 | 4 | 0 | 1 | 0 | 0 | 0 | 0 | 5 |
| ENSLAFG00000028692 | 0 | 1 | 0 | 5 | 0 | 0 | 0 | 0 | 0 | 1 | 15 |
| ENSLAFG00000028299** | 0 | 0 | 0 | 0 | 0 | 0 | 0 | 0 | 0 | 0 | 20 |

*Functional homolog

**”p53 retrogene 9” from Abegglen et al. (2015)
